# Superior Colliculus to VTA pathway controls orienting response to conspecific stimuli

**DOI:** 10.1101/735340

**Authors:** Clément Prévost-Solié, Alessandro Contestabile, Pedro Espinosa, Stefano Musardo, Sebastiano Bariselli, Chieko Huber, Alan Carleton, Camilla Bellone

## Abstract

Social behaviours characterize cooperative, mutualistic aggressive or parental interactions that occurs among conspecifics. Although several neuronal substrates of social behaviour have been identified, whether defined circuits are dedicated to specific aspect of conspecific interaction is still an open question. Ventral Tegmental Area (VTA) contributes to the rewarding properties of conspecific interaction. However, how information related to conspecifics are conveyed to the VTA is still largely unknown. In this study, we identified a population of Superior Colliculus (SC) neurons projecting to the VTA which increase their activity before conspecific interaction and control orienting response towards unfamiliar conspecifics. Finally, we show that SC inputs target a subpopulation of Dopamine (DA) neurons within the VTA that in turn project to dorsolateral striatum (DLS). Our work supports the hypothesis that specialized sub-circuits are dedicated to process different aspect of social interaction.

## Introduction

The term “social interaction” describes interactions with conspecifics, which are highly complex behaviours guided by environmental and internal states. When entering in a novel setting, individuals need to constantly integrate external sensory cues with internal states to properly orient towards conspecifics. These processes prepare the subject to deal with the different situations that a stimulus may evoke *(1)*. At a biological level, several brain regions are functionally relevant for conspecific interaction *(1)*. Therefore, to fully understand the complexity of this behaviour, the neurocircuitry underlying specific aspects of social interaction needs to be precisely investigated.

Because of its role in reward and motivation, the ventral tegmental area (VTA) and dopamine (DA) have long been implicated in conspecific interactions *(2, 3)*. Indeed, DA release peaks upon initial contact with a conspecific and habituates upon subsequent presentation of the same stimulus *(3)*. More recently, it has been shown that the activity of DA neurons encode key aspects of social interaction *(4)*. Interestingly, while stimulation of VTA DA neurons projecting to the NAc (VTA^DA^ – NAc) increases interaction *(4)*, chemogenetic inhibition of VTA DA neurons attenuates both the exploration and the reinforcing properties of novel conspecifics in mice *(5)*. However, the circuit mechanisms that modulate the activity of DA neurons during the interaction with novel conspecifics are still largely unknown.

Novel stimuli, according to Sokolov, generate orienting responses which depend upon the significance of the stimulus and habituate with stimulus repetition *(6)*. Importantly, Pavlov stressed that “orienting responses are necessary in term of organism’s survival. If it were absent, the life of the animal would be in constant danger”. Orienting response represents therefore a complex reaction to a significant event and is the basis of natural selective attention *(7)*. In this framework, social cues are environmental salient stimuli that capture attention and can generate orienting response. After orienting response towards a conspecific is generated, the individual evaluates the situation and decides how to adapt his behaviour and whether to initiate or to avoid the interaction. Orienting response may therefore represent the first and necessary step to engage in social behaviour. Which neural circuits control orienting responses, especially toward conspecific, and how they affect social interaction are still open questions.

Unexpected biologically-relevant salient events evoke DA neuron responses in several species *(8*, *9)*. Indeed, midbrain DA neurons respond with short-latency to different sensory cues *(10)* and their response habituate rapidly when the stimulus is repeated. Thereby, these data suggest that VTA DA neurons could play a role in orienting response toward conspecific during social interaction. Several studies have provided evidence that a subcortical structure in the midbrain, the Superior Colliculus (SC), is a source of short-latency visual inputs to midbrain DA neurons *(11*, *12, 13*, *14)*. SC is an evolutionary conserved midbrain structure that is activated by biological salient stimuli independently of their valence *(15)* and plays a key role in controlling orientation and spatial attention *(16,*, *17, 18)*. Recently, it has been shown that SC sends excitatory projections onto VTA GABA neurons to control defensive behavior while SC inputs onto VTA DA neurons might contribute to reinforcement learning *(19)*. However, whether SC conveys information to VTA DA neurons during social orienting, thus having a pivotal role in conspecific interaction, remains unknown.

Midbrain DA neurons cluster in different subpopulations that regulate emotional, cognitive and motor functions. Efferent projections mainly target the striatum and the cortex and several afferent projections have been anatomically identified. Interestingly, it is generally assumed that DA neurons in the VTA project to the ventral part of the striatum, while DA neurons in the substantia nigra (SN) send projection to the dorsal striatum. However, a gradient distribution of DA cells projecting to the ventromedial, central and dorsolateral striatum has been observed *(20)*. Through a spiral input – output connection between the striatum and the midbrain, information flows from limbic to motor circuits providing a mechanism by which motivation can influence decision-making and action performance *(21)*. Thus, this cross-talk between circuits would be important to adjust behaviour in function of novel information. Whether different loops within the midbrain circuit play a different role in conspecific interaction is still largely unknown.

Here we show that the neuronal activity of identified SC neurons projecting to VTA (SC – VTA) increases before nose-to-nose interaction between two unfamiliar conspecifics and during orienting response toward unfamiliar conspecifics. Optogenetic manipulation of SC – VTA pathway disturbs orienting response towards social stimuli and bilaterally interfere with conspecific interaction. Interestingly, the activity of medial Prefrontal cortex (mPFC) to VTA input pathway (mPFC – VTA) increases during conspecific interaction but not during orienting response toward the social stimulus. Remarkably, we found that DA neurons receiving SC projections are located in the lateral VTA and mainly provide inputs to dorsolateral striatum (DLS). Finally, optogenetic stimulation of the VTA^DA^ – DLS pathway affects conspecific interaction similarly to SC – VTA stimulation.

## Results

### VTA receives projection from SC

To anatomically assess projections from SC to VTA, we first injected an adeno-associated virus (AAV) expressing a yellow fluorescent protein (eYFP; AAV5-hSyn-eYFP) in the SC (**Figures 1A-B**), and we observed SC axons in the VTA (**Figure 1B’**). Then, to identify the anatomical position of SC neurons projecting to VTA, we injected a retrograde virus (CAV) expressing Cre in the VTA and a Cre-dependent AAV virus expressing mCherry (AAV5-hSyn-DIO-mCherry) in the SC (**Fig. 1C**). This experiment revealed that SC neurons projecting to the VTA are predominantly located in the intermediate and deep layers (**Figures 1D** and **F**). Furthermore, immunohistochemistry analysis indicated that 88,1% of SC neurons projecting to the VTA are CaMKII positive (**Figures. 1E-F**), suggesting a prevalent excitatory connection between SC and VTA. To explore the functional connectivity of SC – VTA projections, we first injected an AAV5-hSyn-ChR2-eYFP in SC and performed whole-cell patch clamp recordings to confirm that the SC neurons followed light stimulation protocol (**Figure 1G**). In a second time, we infected mice expressing the Cre under the promoter of the DA transporter (DAT-Cre) or under the glutamic acid decarboxylase (GAD65-Cre) with an AAV-DIO-mCherry in VTA to target respectively VTA DAT^+^ and GAD^+^ neurons. The mice were subsequently injected with AAV5-hSyn-ChR2-eYFP in SC (**Figure 1H**) to explore the functional connectivity between SC and VTA DAT^+^ and GAD^+^ neurons. Interestingly, optogenetic stimulation of axons in the VTA evoked excitatory postsynaptic currents (EPSCs) and revealed connectivity both for VTA DAT^+^ (25.7%) and GAD^+^ (42.3%) neurons (**Figure 1I**). The delay to induce a response through the optogenetic stimulation confirmed that the connection was monosynaptic (**Figure 1J**). Remarkably, the percentage of evoked EPSC was higher than IPSC for VTA DAT^+^ cells (25.7% vs 4.4%) strengthening the prevalent excitatory connection between SC and VTA (**Sup. Figure 1A-C**). Interestingly, mapping a part of the VTA cells connected to the SC, we observed that DAT^+^ connected neurons were located in the lateral part of the VTA while GAD^+^ connected neurons were located in medial VTA (**Figure 1K-K’**). Taken together, these results show that distinct neuronal populations within the VTA receive inputs from SC.

**Figure 1:**
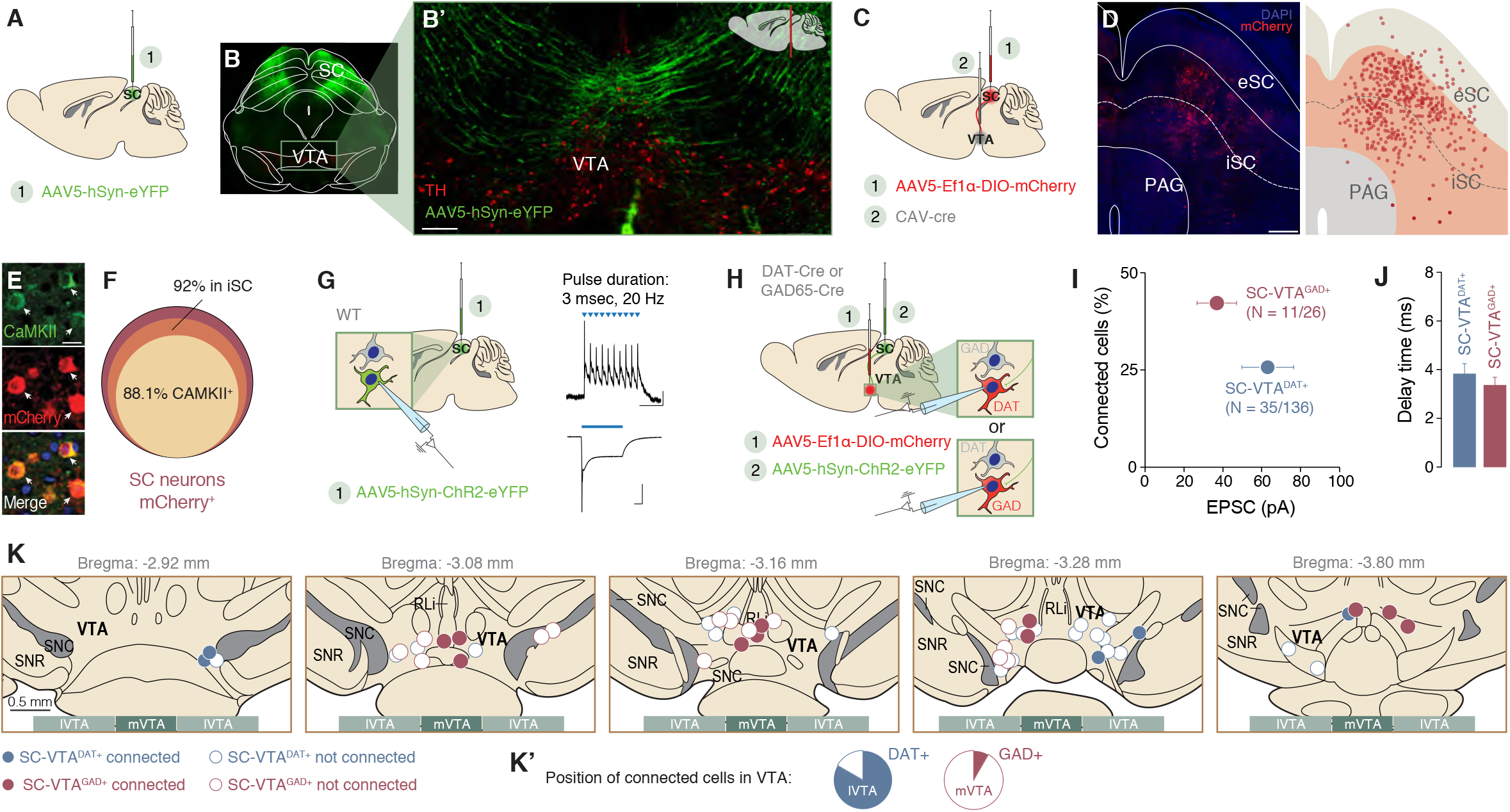
Anatomo-functional connectivity from SC to VTA dopaminergic and GABAergic neurons. **(A)** Schema of injection in the Superior Colliculus (SC) with AAV5-hSyn-eYFP. **(B)** Representative coronal image of immuno-staining experiments against Tyrosine Hydroxylase (TH) enzyme (in red) performed on midbrain slices of adult mice infected with AAV5-hSyn-eYFP (green) in the SC. The projecting fibers from SC to VTA are visible. **(B’)** Image at higher magnification of the coronal slice (scale bar: 100 *μ*m). The fibers project from SC to VTA. **(C)** Schema of injections in the VTA with CAV-Cre and AAV5-Ef1α-DIO-mCherry in the SC. **(D)** Left: Representative image of the infected cells with CAV-Cre and AAV5-Ef1α-DIO-mCherry in the SC (scale bar: 200 *μ*m). Right: schema reporting the position of mCherry positive cells in the SC for 4 infected brains. **(E)** Representative image of immuno-staining against Ca^2+^/calmodulin-dependent protein kinase II (CAMKII) in the SC and infected cells with AAV5-Ef1α-DIO-mCherry (scale bar: 10 *μ*m). **(F)** Quantification of infected cells in the SC. The cells are preferentially located in the intermediate layer of the SC and are predominantly CAMKII^+^. **(G)** Whole cell patch clamp from SC ChR2-expressing neurons. Protocol of stimulation indicates that SC neurons follow 20 Hz light stimulation protocol. **(H)** Schema of injection in the SC with AAV5-hSyn-ChR2-eYFP and patch of the VTA DAT^+^ and GAD^+^ neurons (depending on the mice line). **(I)** Quantification of connected cells from SC onto VTA DAT^+^ and GAD^+^ neurons in relation of the amplitude of EPSC. **(J)** Delay time of the EPSC for VTA DAT^+^ and GAD^+^ neurons. **(K-K’)** Position of some patched VTA DAT^+^ and GAD^+^ neurons. SC-VTA^DAT+^ connected neurons are mainly in the lateral part of the VTA (lVTA) while SC-VTA^GAD+^ connected neurons are more medially located (mVTA). N indicates number of cells and error bars report s.e.m.

### SC – VTA neuron activity increases before social interaction

Both SC and VTA have been previously implicated in social behaviour *(22) (4)* (Prévost-Solié, Girard et al., 2020) but evidences of the involvement of SC – VTA pathway in social interaction are still missing. To investigate the roles of SC – VTA in social interaction, we targeted the SC neurons projecting to the VTA by injecting a retrograde AAV encoding Cre virus (AAVrg-Ef1α-Cre-mCherry) in the VTA and a Cre-dependent AAV encoding GCaMP6s in the SC (AAV9-FLEX-GCamp6s; **Figures 2A-B’**). We then implanted the optic fiber in the SC, and recorded the Ca^2+^ transient during free social interaction with a juvenile sex matched conspecific (**Figures 2C**). By time-locking the Ca^2+^ transient on specific events, we observed an increase in SC – VTA pathway activity starting before nose-to-nose (**Figures 2D-E**) and nose-to-body contact (**Figures 2F-G**). On the other hand, no changes were observed before or after passive contacts with the conspecific (**Figures 2H-I**) or rearing episodes (**Figures 2J-K**). Taken together, these results show that the activity of the SC population that project to VTA increases before active contacts during free and direct social interaction task.

**Figure 2:**
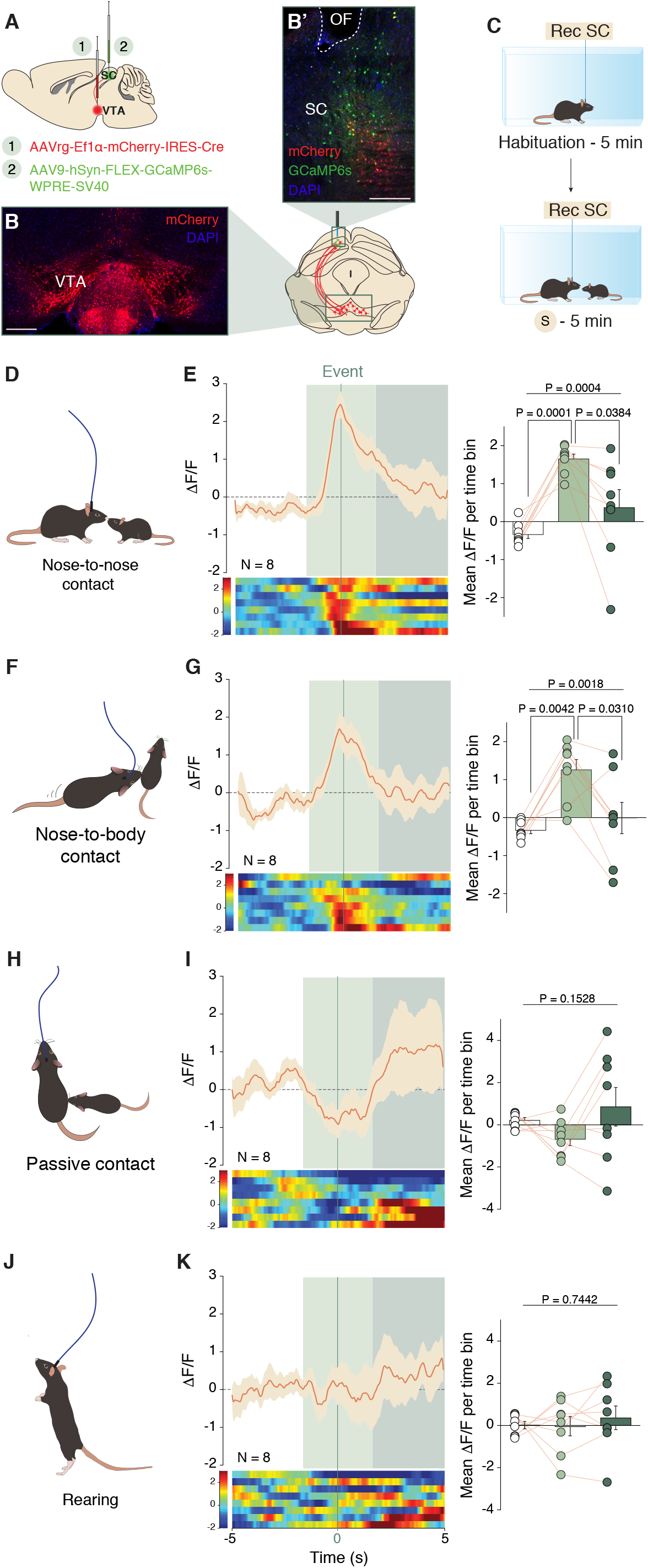
Calcium activity of SC to VTA projecting neurons during free social interaction. **(A)** Schema of injections of AAVrg-Ef1α-mCherry-IRES-Cre in the VTA and AAV9-hSyn-FLEX-GCaMP6s-WPRE-SV40 in the SC. Representative coronal images of midbrain slices of adult mice infected with AAVrg-Ef1α-mCherry-IRES-Cre in the VTA **(B,** scale bar: 100 *μ*m) and AAV9-hSyn-FLEX-GCaMP6s-WPRE-SV40 in the SC **(B’,** scale bar: 50 *μ*m). **(C)** Schema of free social interaction task. **(D, F, H** and **J)** Time locked events observed during free social interaction. **(E)** Left: Peri-event time histogram (PETH) of normalized calcium activity of SC – VTA projecting neurons, centered on nose-to-nose contacts. Right: Normalized calcium activity (Z-score) before during and after nose-to-nose. RM one-way ANOVA (Events main effect: F_(2,7)_ = 10.3, P = 0.0018) followed by Bonferroni-Holm post-hoc test correction. **(G)** Left: PETH of normalized calcium activity of SC – VTA projecting neurons, centered on nose-to-body contacts. Right: Normalized calcium activity (Z-score) before during and after nose-to-body. RM one-way ANOVA (Events main effect: F_(2,7)_ = 14.15, P = 0.0004) followed by Bonferroni-Holm post-hoc test correction. **(I)** Left: PETH of normalized calcium activity of SC – VTA projecting neurons, centered on passive contacts. Right: Normalized calcium activity (Z-score) before during and after passive contacts. RM one-way ANOVA (Events main effect: F_(2,7)_ = 2.15, P = 0.1528). **(K)** Left: PETH of normalized calcium activity of SC – VTA projecting neurons, centered on rearing behaviour. Right: Normalized calcium activity (Z-score) before during and after rearing. RM one-way ANOVA (Events main effect: F_(2,7)_ = 0.3017, P = 0.7442).N indicates number of mice. Error bars report s.e.m.

### SC – VTA pathway bidirectionally controls social interaction

To test the role of SC – VTA activity during social interaction, we injected blue-light sensitive ChR2 (AAV-hSyn-ChR2-eYFP or AAV-hSyn-eYFP as control) or red light-sensitive optogenetic inhibitor Jaws (AAV-hSyn-Jaws-GFP or AAV-hSyn-eYFP as control) in the SC. In a second time, we implanted an optic fiber over the VTA to stimulate or inhibit SC axon terminals (**Figures 3A-C’**). As control, we tested that Jaws activation induces terminal inhibition by recording light induced EPSCs in the VTA after injection of Jaws/ChR2 in the SC (**Sup. Figure 2A-C**). We then placed the mice in a home cage-like arena and, after 3 min of baseline, we exposed the experimental mouse to an unfamiliar conspecific for 2 min (**Figure 3C**). During the entire light ON condition (5 min), mice were photostimulated or photoinhibited according to the protocols used (see **Figure 3C’** for stimulation/inhibition protocols). The experiment was then repeated with another unfamiliar conspecific after 3h delay (counterbalanced), and we compared the total time spent in investigation (time sniffing) between light OFF and light ON epochs. Interestingly, the photostimulation of SC – VTA pathway in ChR2-expressing mice decreased time interaction without affecting the eYFP-control group (**Figure 3D**). The time-course of the interaction with the unfamiliar conspecific revealed that the eYFP-expressing mice quickly habituate to the stimulus and spend less time investigating the conspecific through the 2 min of interaction (**Figure 3E**). Interestingly, ChR2-expressing mice decreased their interaction time especially over the first 30 seconds when comparing light ON and OFF conditions (**Figure 3E**). Contrarily to photostimulation, photoinhibition of this pathway significantly increased investigation time in Jaws-expressing mice without affecting the eYFP-control group (**Figures 3F-G**). Furthermore, to better dissect the behavioural effects of SC – VTA pathway manipulation, we analysed detailed aspects of social and non-social behaviour sequences during both light OFF and light ON epochs (**Sup. Figure 3A**). Although we did not find differences in rearing behaviour, photostimulation induced less following and nose-to-nose interaction (**Sup. Figures 3B-D**), while photoinhibition elicited opposite changes only in following behaviour between light ON and OFF conditions **(Sup. Figures 3E-G)**. These data show that SC – VTA pathway bidirectionally controls free social interaction.

**Figure 3:**
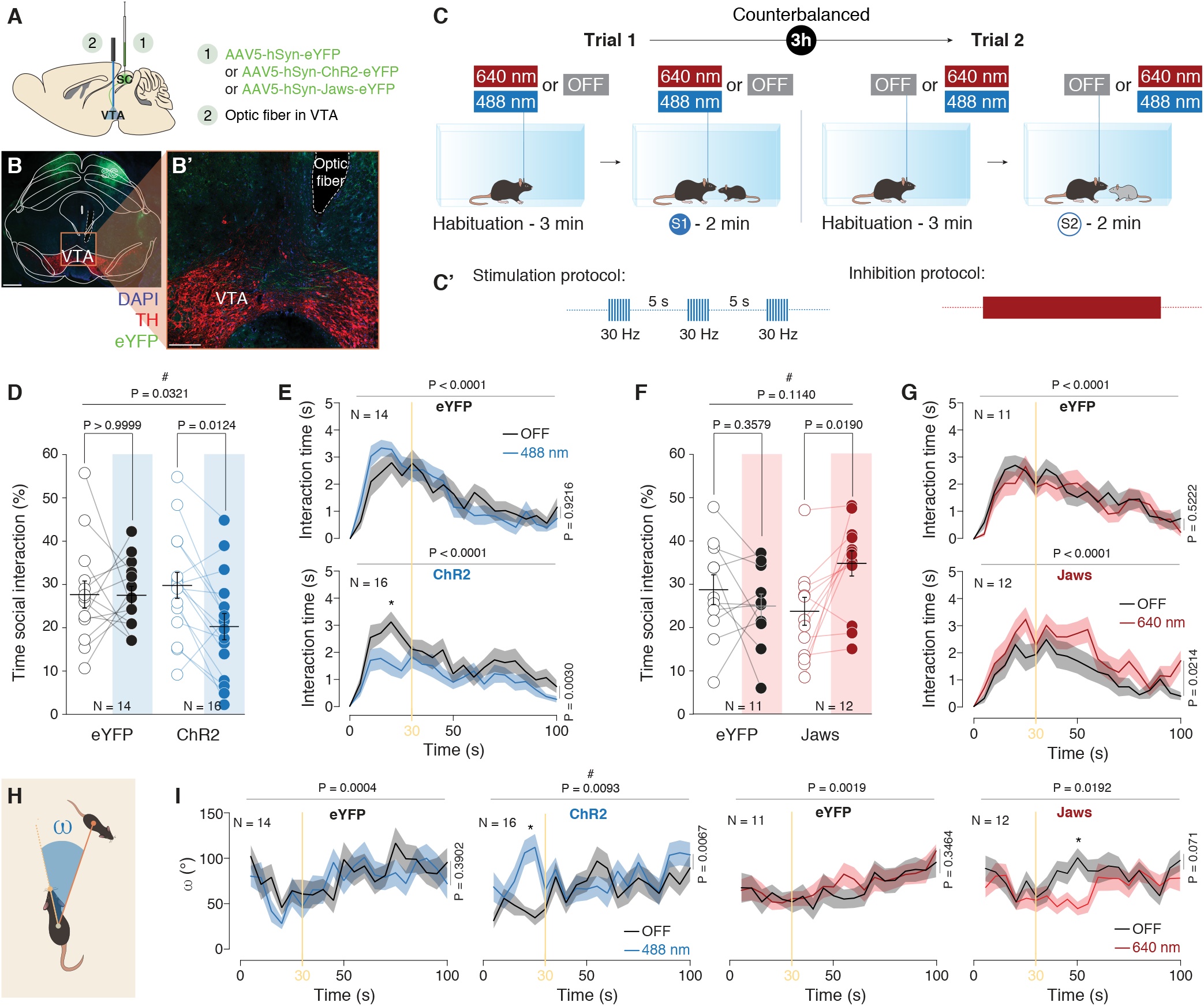
Optogenetic manipulation of SC – VTA pathway alters social interaction and head orientation towards conspecific. **(A)** Schema of injections sites in SC with AAV5-hSyn-eYFP, AAV5-hSyn-ChR2-eYFP or AAV5-hSyn-Jaws-GFP, and optic fiber implantation above the VTA. **(B**) Representative image of coronal midbrain slices of adult mice infected with AAV5-hSyn-eYFP (green) in the SC. In red is visible the immunostaining anti-Tyrosine Hydroxylase (TH) (scale bar: 500 μm). **(B’)** Image at higher magnification of the VTA of B (scale bar: 100 *μ*m). **(C)** Schema of free social interaction. The eYFP-, ChR2- and Jaws-expressing mice freely interacted with two different unfamiliar mice under both stimulation conditions. **(C’)** Stimulation and inhibition protocols. 8 pulses of 488 nm light (30 Hz) were separated by 5s in the light ON condition. Continuous inhibition was instead provoked with 640 nm light. **(D)** Time social interaction during the free social interaction task for eYFP- and ChR2-expressing mice in the SC. RM two-way ANOVA (Light main effect: F_(1,28)_ = 5.0855, P = 0.0321; Virus main effect: F_(1,28)_ = 0.8528, P = 0.3637; Light x Virus Interaction: F_(1,28)_ = 4.1962, P = 0.0500) followed by Bonferroni-Holm post-hoc test correction. **(E)** Time course of the time of social interaction for eYFP- and ChR2-expressing mice in the SC in light and no-light stimulation conditions. RM two-way ANOVA (eYFP: Light main effect: F_(1,13)_ = 0.01008, P = 0.9216; Time main effect: F_(20,260)_ = 13.52, P < 0.0001; Light x Time Interaction: F_(20,260)_ = 1.047, P = 0.4074. ChR2: Light main effect: F_(1,15)_ = 12.46, P = 0.0030; Time main effect: F_(20,300)_ = 10.35, P < 0.0001; Light x Time Interaction: F_(20,300)_ = 0.7968, P = 0.7173) followed by Bonferroni’s multiple comparisons post-hoc test. **(F)** Time social interaction during the free social interaction task for eYFP- and Jaws-expressing mice in the SC. RM two-way ANOVA (Light main effect: F_(1,21)_ = 2.7201, P = 0.1140; Virus main effect: F_(1,21)_ = 0.1985, P = 0.6605; Light x Virus Interaction: : F_(1,21)_ = 8.3378, P = 0.0088) followed by Bonferroni-Holm post-hoc test correction. **(G)** Time course of the time of social interaction for eYFP- and Jaws-expressing mice in the SC in light and no-light stimulation conditions. RM two-way ANOVA (eYFP: Light main effect: F_(1,10)_ = 0.4399, P = 0.5222; Time main effect: F_(20,200)_ = 7.902, P < 0.0001; Light x Time Interaction: F_(20,200)_ = 0.5527, P = 0.9398. Jaws: Light main effect: F_(1,11)_ = 7.188, P = 0.0214; Time main effect: F_(20,220)_ = 7.509, P < 0.0001; Light x Time Interaction: F_(20,220)_ = 0.6139, P = 0.9005) followed by Bonferroni’s multiple comparisons post-hoc test. **(H)** Schema representing the points and vectors used for the calculation of the oriented angle towards the social stimulus (ω). **(I)** Time course of ω for eYFP-, ChR2- and Jaws-expressing mice in the SC during free social interaction task in light and no-light stimulation conditions. RM two-way ANOVA (eYFP (OFF - 488 nm): Light main effect: F_(1,13)_ = 0.7902, P = 0.3902; Time main effect: F_(20,260)_ = 5.513, P < 0.0001; Light x Time Interaction: F_(20,260)_ = 0.8264, P = 0.6806. ChR2 (OFF - 488 nm): Light main effect: F_(1,15)_ = 10.38, P = 0.0067; Time main effect: F_(20,300)_ = 4.838, P < 0.0001; Light x Time Interaction: F_(20,300)_ = 0.2049, P = 0.0060. eYFP (OFF - 640 nm): Light main effect: F_(1,10)_ = 0.9762, P = 0.3464; Time main effect: F_(19,190)_ = 2.349, P = 0.0019; Light x Time Interaction: F_(19,190)_ = 0.3207, P = 0.9972. Jaws (OFF - 640 nm): Light main effect: F_(1,11)_ = 3.998, P = 0.0709; Time main effect: F_(19,209)_ = 1.853, P = 0.0192; Light x Time Interaction: F_(19,209)_ = 1.14, P = 0.3136) followed by Bonferroni’s multiple comparisons post-hoc test. N indicates number of mice. # indicates significantly different interaction. Error bars report s.e.m.

When an unfamiliar conspecific is introduced in the environment, the experimental mouse typically stops the ongoing action to re-orient its attention towards the stimulus, right before or while initiating the interaction. The SC is highly involved in attention and orienting response toward novel and salient stimuli *(16) (17) (18)* and our previous results with the fiber photometry suggested that the SC – VTA neuron activity occurs before the interaction. Thereby, to capture this attentional shift during social exposure, we calculated in ChR2 and Jaws-expressing mice the head orientation (ω) as the angle formed between the vector determined from the body center to the nose of the experimental mouse, to the juvenile body center (**Figure 3H**). In eYFP-control mice, the angle of head orientation follows the interaction time and rapidly increases during the first 100 seconds of the task, revealing reduced orientation towards the unfamiliar conspecific across the session (**Figure 3I**). This result reflects an increased attention directed towards the unfamiliar conspecific during the first instants of social interaction. Interestingly, photostimulation of SC – VTA pathway decreases orientation towards the social stimulus while photoinhibition prolongs this response in the first half of the test (**Figure 3I**). Thereby, photoinhibition and photostimulation of SC – VTA pathway alters the orientation compared to light OFF epochs in opposite ways. Remarkably, we noticed a significant inversed linear correlation across the session between interaction time and head-orientation angle for eYFP-control and Jaws-expressing animals during both light OFF and light ON epochs (**Sup. Fig. 4A-B**). On the other hand, we observed a loss of correlation during light ON epochs in ChR2-expressing mice only (**Sup. Fig. 4A)**. These results suggest that orientation and social contact are tightly related features of social behaviour and that SC – VTA pathway may influence social interaction by encoding orienting response.

**Figure 4:**
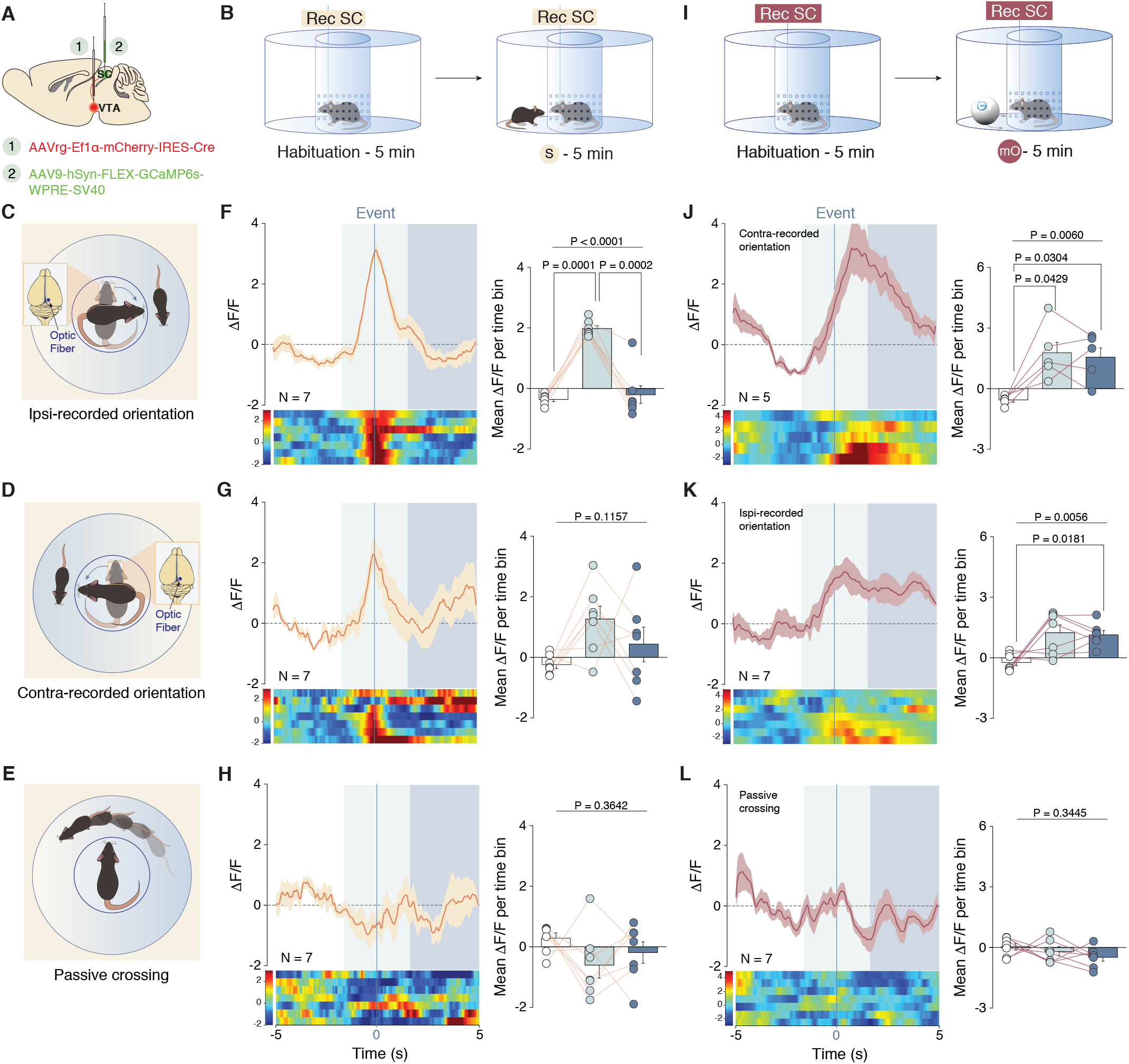
Calcium activity of SC to VTA projecting neurons during social and non-social orientation task. **(A)** Schema of injections of AAVrg-Ef1α-mCherry-IRES-Cre in the VTA and AAV9-hSyn-FLEX-GCaMP6s-WPRE-SV40 in the SC. **(B)** Schema of the social orientation task. **(C**, **D** and **E)** Time locked events observed during social orientation task. **(F)** Left: Peri-event time histogram (PETH) of normalized calcium activity of SC – VTA projecting neurons, centered on ipsi-recorded orientation for social stimulus. Right: Normalized calcium activity (Z-score) before during and after ipsi-recorded orientation. RM one-way ANOVA (Events main effect: F_(2,6)_ = 44.34, P < 0.0001) followed by Bonferroni-Holm post-hoc test correction. **(G)** Left: PETH of normalized calcium activity of SC – VTA projecting neurons, centered on contra-recorded orientation. Right: Normalized calcium activity (Z-score) before during and after contra-recorded orientation. RM one-way ANOVA (Events main effect: F_(2,6)_ = 2.59, P = 0.1157). **(H)** Left: PETH of normalized calcium activity of SC – VTA projecting neurons, centered on passive crossing. Right: Normalized calcium activity (Z-score) before during and after passive crossing. RM one-way ANOVA (Events main effect: F_(2,6)_ = 1.1002, P = 0.2611). **(I)** Schema of non-social orientation task with a moving ball (mO). **(J)** Left: Peri-event time histogram (PETH) of normalized calcium activity of SC – VTA projecting neurons, centered on ipsi-recorded orientation for moving object stimulus. Right: Normalized calcium activity (Z-score) before during and after ipsi-recorded orientation. RM one-way ANOVA (Events main effect: F_(2,4)_ = 10.34, P = 0.0060) followed by Bonferroni-Holm post-hoc test correction. **(K)** Left: PETH of normalized calcium activity of SC – VTA projecting neurons, centered on contra-recorded orientation. Right: Normalized calcium activity (Z-score) before during and after contra-recorded orientation. RM one-way ANOVA (Events main effect: F_(2,6)_ = 8.06, P = 0.0056) followed by Bonferroni-Holm post-hoc test correction. **(L)** Left: PETH of normalized calcium activity of SC – VTA projecting neurons, centered on passive crossing. Right: Normalized calcium activity (Z-score) before during and after passive crossing. RM one-way ANOVA (Events main effect: F_(2,6)_ = 1.17, P = 0.3445). N indicates number of mice. Error bars report s.e.m.

### SC – VTA neurons are activated during orientation task

Based on the results, we next aimed to better dissect the SC – VTA pathway during orienting response toward conspecific. We therefore designed an orientation task and recorded fiber photometry signals from SC projecting to the VTA. As described previously, we injected an AAVrg-Ef1α-Cre-mCherry in the VTA and an AAV9-FLEX-GCamp6s in the SC **(Figure 4A**). We introduced and restrained the experimental mice in an enclosure positioned in the center of a circular arena allowing it only to turn either on the right or on the left **(Figure 4B**). After 5 min habituation, we placed a sex matched juvenile conspecific in the arena and we recorded the Ca^2+^ transient during ipsilateral **(Figure 4C**) or contralateral (**Figure 4D**) orienting response toward conspecific or during passive crossing (**Figure 4E**). Interestingly, we observed a significant increase in Ca^2+^ transient during ipsi-recorded orienting response (**Figure 4F**) but not during contra-recorded or passive crossing events (**Figure 4G-H**). These data strongly indicate that SC – VTA pathway is highly involved in orienting responses toward the conspecific.

To test if the SC – VTA pathway was also involved in orienting response toward salient novel stimuli, we used a moving object and recorded Ca^2+^ transient in the SC projecting to VTA neurons in the same orientation task (**Figure 4I**). The object was programmed to move similarly to a conspecific, with approximately the same speed. Remarkably, ipsi- and contra-recorded orienting response towards the moving object induced increase in calcium recording while passive crossing did not induced changes in Ca^2+^ transient (**Figure 4J-L**). These data indicate that the SC – VTA pathway encodes the orienting response towards salient stimuli.

### SC – VTA pathway controls orienting response toward conspecific

To further assess the role of SC – VTA pathway in the orientation task, we injected eYFP, ChR2 or Jaws in the SC and placed the fiber optic in the VTA (**Figure 5A**). We stimulated or inhibited the terminals during the task through habituation and orienting response toward conspecific stimuli. The day after we counterbalanced the mice (**Figure 5B-B’**). Analysing the relative position of the social stimulus in respect of the experimental mouse (**Figure 5C**), we observed that in the light ON condition, SC – VTA ChR2-expressing mice passed less time oriented towards the stimulus during the first minute of the task (**Fig. 5D**). Vectorial analysis of the head orientation angle (ω, **Figure 5E**) revealed a habituation in time spent orienting toward the conspecific for the eYFP-expressing mice during the task (**Figure 5F**). On the other hand, SC – VTA ChR2-expressing mice decreased the time orienting toward the stimulus during the first minute and do not habituate within the task (**Figure 5F**). Interestingly, compared to the control condition, Jaws-expressing mice during light ON epoch do not habituate and a remarkable percentage of them increased the orientation time towards the social stimulus during the second minute (**Figure 5F**). All these data strongly support our previous results and highlight the important role of the SC – VTA pathway in orienting response towards salient stimuli as conspecific.

**Figure 5:**
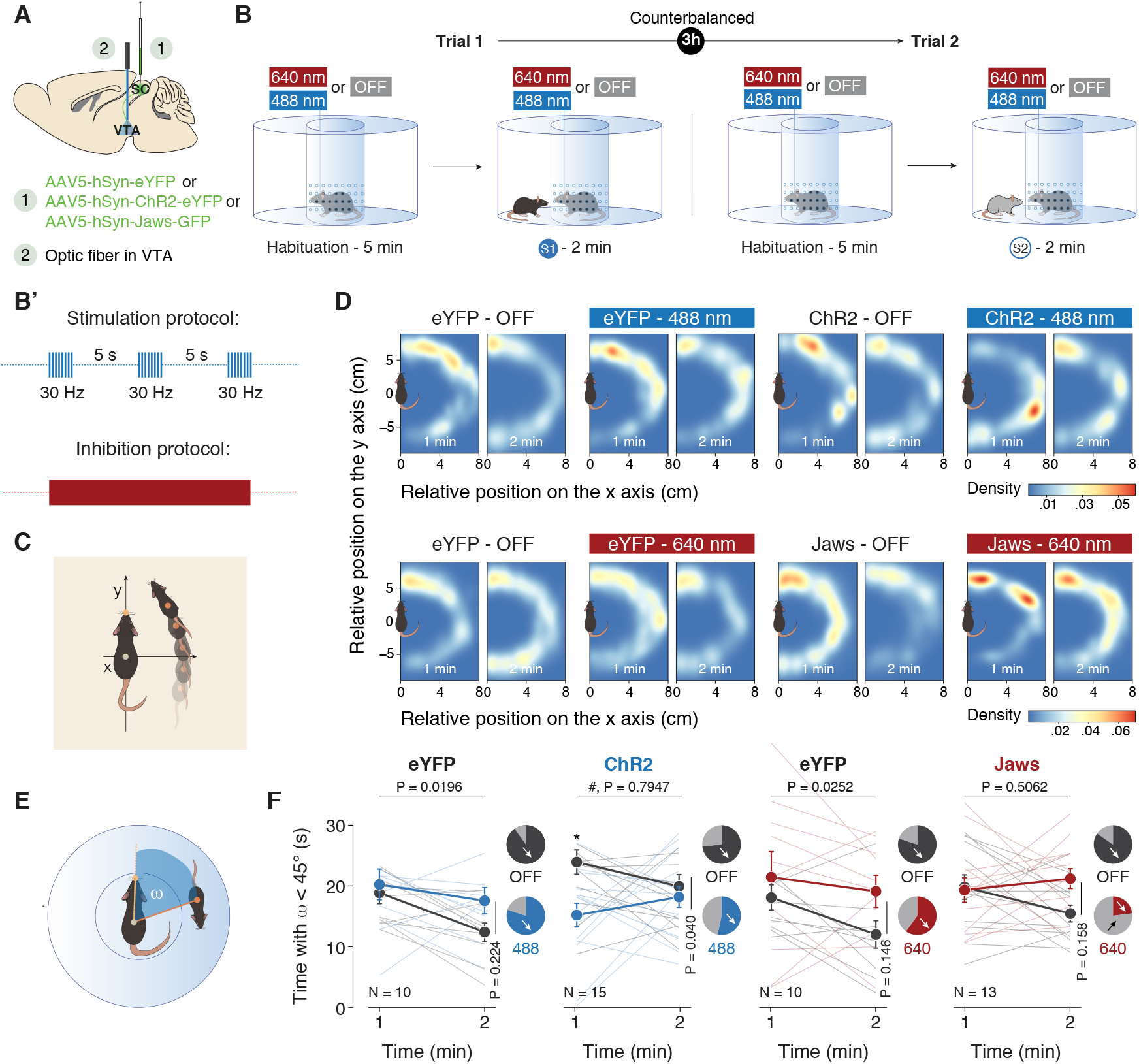
Optogenetic manipulation of SC – VTA pathway alters orienting response. **(A)** Schema of injections sites in SC with AAV5-hSyn-eYFP, AAV5-hSyn-ChR2-eYFP or AAV5-hSyn-Jaws-GFP, and optic fiber implantation above the VTA. **(B)** Schema of the social orientation task. The eYFP-, ChR2- and Jaws-expressing mice oriented towards two different unfamiliar mice under both stimulation conditions. **(B’)** Stimulation and inhibition protocol. Eight pulses of 488 nm light (30 Hz) were separated by 5s in the light ON condition. Continuous inhibition was instead provoked with 640 nm light. **(C)** Schema representing the relative position of the social stimulus when the center body point of the experimental animals is fixed at (0, 0) and the nose point is fixed along the y axis. **(D)** Heatmaps reporting the relative position of the social stimulus during orientation task for the first and second minute in the different conditions. **(E)** Schema representing the points and vectors used for the calculation of the oriented angle towards the social stimulus (ω). **(F)** Time passed with a ω < 45° for the 1^st^ and 2^nd^ minute of the social orienting task in light and no-light conditions. RM two-way ANOVA (eYFP (OFF - 488 nm): Light main effect: F_(1,10)_ = 1.683, P = 0.2236; Time main effect: F_(1,10)_ = 7.711, P = 0.0196; Light x Time Interaction: F_(1,10)_ = 1.254, P = 0.2890. ChR2 (OFF – 488 nm): Light main effect: F_(1,14)_ = 5.138, P = 0.0398; Time main effect: F_(1,14)_ = 0.070, P = 0.7947; Light x Time Interaction: F_(1,14)_ = 4.868, P = 0.0446. eYFP (OFF - 640 nm): Light main effect: F_(1,9)_ = 2.537, P = 0.1456; Time main effect: F_(1,9)_ = 7.185, P = 0.0252; Light x Time Interaction: F_(1,9)_ = 0.6144, P = 0.4533. Jaws (OFF - 640 nm): Light main effect: F_(1,12)_ = 2.266, P = 0.1581; Time main effect: F_(1,12)_ = 0.4696, P = 0.5062; Light x Time Interaction: F_(1,12)_ = 4.572, P = 0.0538) followed by Bonferroni’s multiple comparisons post-hoc test. Pie charts represent the percentage of mouse that decrease (D) the orientation between 1^st^ and 2^nd^ minute. N indicates number of mice. # indicates significantly different interaction. Error bars report s.e.m.

### SC – VTA pathway stimulation increases exploratory behaviour and does not induce place preference

Finally, to test whether manipulation of SC-VTA pathway changes exploratory behaviour non-associated with a stimulus, eYFP-, ChR2- or Jaws-expressing mice were placed in an open field for 10 min to assess their locomotor activity (**Sup. Figures 5A**). Although we did not observe significant differences between light OFF and light ON epochs using a within-group analysis, ChR2-expressing mice increased their distance moved relative to control animals upon SC terminal stimulation (**Sup. Figure 5B-D**). On the other hand, no changes in distance moved where observed in Jaws-expressing mice between light OFF and light ON epochs (**Sup. Figure 5E-G**). To exclude anxiolytic effects due to SC – VTA pathway alteration, we measured the time spent in the center of the arena, which was similar between eYFP-control, ChR2-expressing and Jaws-expressing mice (**Sup. Figure 5H-I**). We finally tested whether manipulation of SC – VTA pathway would support real-time place preference. Animals were placed into a two-chambered arena, where only one chamber was paired with optical stimulation (**Sup. Figure 5J**). eYFP-control, ChR2-expressing or Jaws-expressing mice spent a comparable amount of time in each chamber (**Sup. Figure 5K-L**).

These data indicate that SC – VTA pathway plays a specific role in the orientation toward the conspecific during the interaction.

### The mPFC – VTA pathway is differently activated during social behaviour compared to the SC – VTA pathway and is not implicated in orienting response

In order to test the specificity of this pathway in orienting response, we investigated the role of a different source of excitatory inputs to the VTA during social interaction, the medial Prefrontal Cortex (mPFC) (Beier et al. 2015). We targeted the mPFC neurons projecting to the VTA by injecting a retrograde AAV encoding Cre virus (AAVrg-Ef1α-mCherry-IRES-Cre) in the VTA and a Cre-dependent AAV encoding GCaMP6s in the mPFC (AAV9-hSyn-FLEX-GCamp6s-WPRE-SV40; **Figure 6A-B**). We then implanted the optic fiber in the mPFC, and recorded the Ca^2+^ transient during free social interaction with a juvenile sex matched conspecific (**Figure 6C**) and the orientation task (**Figure 6G**). Interestingly, we observed a prolonged increase in Ca^2+^ transient at mPFC – VTA neurons after nose-to-nose (**Figure 6D**) and nose-to-body contact (**Figure 6E**) but not during passive interaction (**Figure 6F**). Remarkably, the kinetic of the calcium signals in the mPFC is significantly different from the kinetic observed in the SC suggesting a different implication on social behaviour of the mPFC – VTA projections. Moreover, no significant activation of the mPFC – VTA pathway was observed during the orientation task (**Figure 6G-M**).

**Figure 6:**
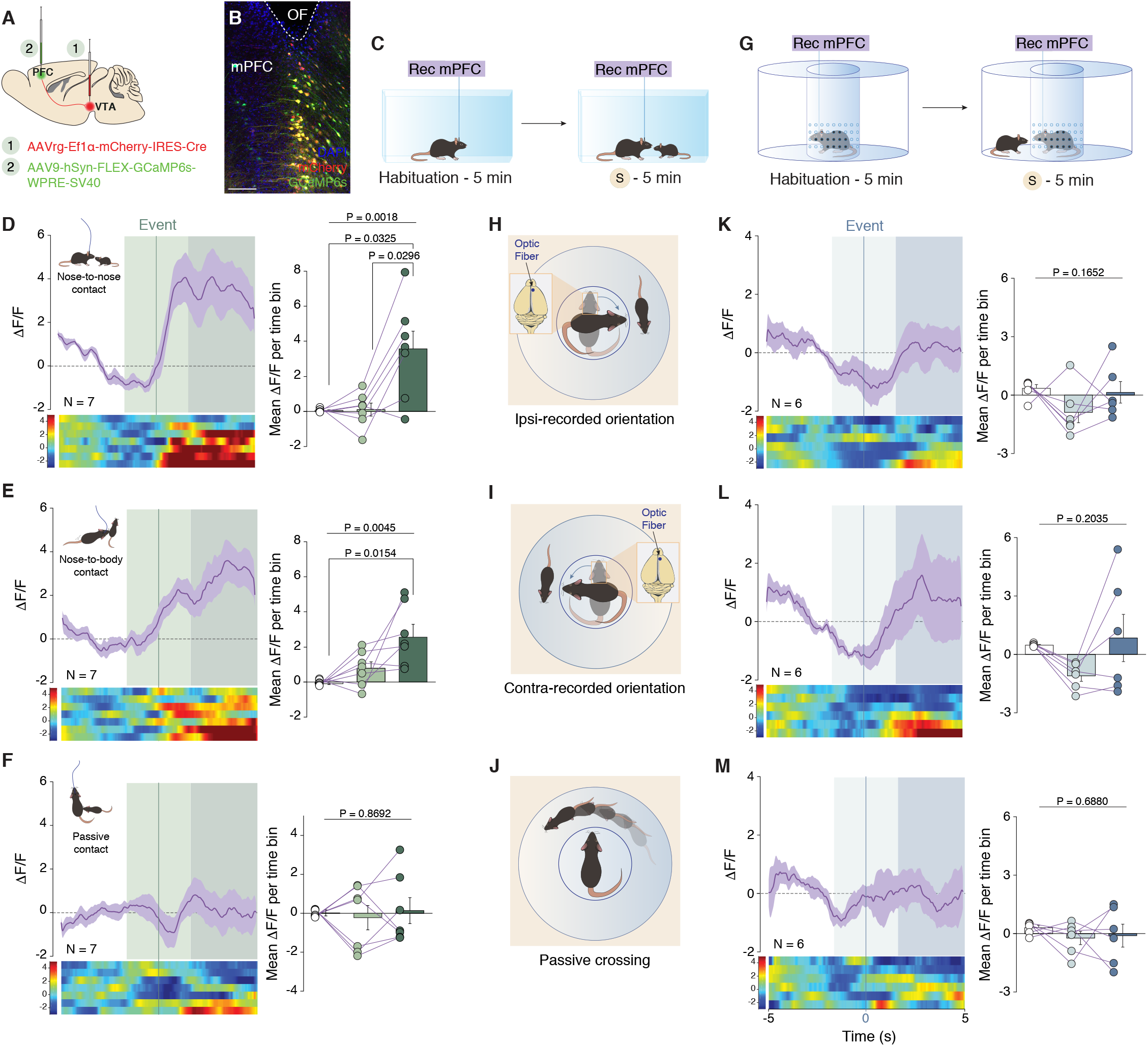
Calcium activity of mPFC – VTA projecting neurons during free social interaction and social orientation task. **(A)** Schema of injections of AAVrg-Ef1α-mCherry-IRES-Cre in the VTA and AAV9-hSyn-FLEX-GCaMP6s-WPRE-SV40 in the mPFC. **(B)** Representative coronal image of mPFC slices of adult mice infected with AAVrg-Ef1α-mCherry-IRES-Cre in the VTA and AAV9-hSyn-FLEX-GCaMP6s-WPRE-SV40 in the mPFC (scale bar: 50 *μ*m). The location of the optic fiber (OF) is indicated. **(C)** Schema of free social interaction. **(D)** Left: PETH of normalized calcium activity of mPFC – VTA projecting neurons, centered on nose-to-nose contacts. Right: Normalized calcium activity (Z-score) before during and after nose-to-nose. RM one-way ANOVA (Events main effect: F_(2,6)_ = 11.28, P = 0.0018) followed by Bonferroni-Holm post-hoc test correction. **(E)** Left: PETH of normalized calcium activity of mPFC – VTA projecting neurons, centered on nose-to-body contacts. Right: Normalized calcium activity (Z-score) before during and after nose-to-body. RM one-way ANOVA (Events main effect: F_(2,6)_ = 8.79, P = 0.0045) followed by Bonferroni-Holm post-hoc test correction. **(F)** Left: PETH of normalized calcium activity of mPFC – VTA projecting neurons, centered on passive contacts. Right: Normalized calcium activity (Z-score) before during and after passive. RM one-way ANOVA (Events main effect: F_(2,6)_ = 0.14, P = 0.8692. **(B)** Schema of the social orientation task. **(H**, **I** and **J)** Time locked events observed during social orientation task. **(K)** Left: PETH of normalized calcium activity of mPFC – VTA projecting neurons, centered on ispsi-recorded orientation. Right: Normalized calcium activity (Z-score) before during and after ipsi-recorded orientation. RM one-way ANOVA (Events main effect: F_(2,5)_ = 2.17, P = 0.1652). **(L)** Left: PETH of normalized calcium activity of mPFC – VTA projecting neurons, centered on contra-recorded orientation. Right: Normalized calcium activity (Z-score) before during and after contra-recorded orientation. RM one-way ANOVA (Events main effect: F_(2,5)_ = 1.87, P = 0.2035). **(M)** Left: PETH of normalized calcium activity of mPFC – VTA projecting neurons, centered on passive crossing. Right: Normalized calcium activity (Z-score) before during and after passive crossing. RM one-way ANOVA (Events main effect: F_(2,5)_ = 0.39, P = 0.6880). N indicates number of mice. Error bars report s.e.m.

Taken together, these results indicate that mPFC – VTA and SC – VTA pathways are activated at different time points during the conspecific interaction suggesting that the two pathways convey different information.

### The SC projects to VTA DAT^+^ neurons connected with DorsoLateral Striatum (DLS)

We next tested the hypothesis that SC targets a neuronal sub-population of the VTA that is anatomically different from the one targeted by the mPFC. Midbrain DAT^+^ neuron axons innervate the striatum with a gradient distribution of cells projecting to the ventromedial central and dorsolateral striatum *(21)*. We therefore injected Cholera Toxin Subunit B (CTB)-488 in the Nucleus Accumbens (NAc) and CTB-555 in the Dorsolateral part of the striatum (DLS) and we imaged the VTA (**Figure 7A-D**). As previously reported *(23)*, we found that VTA neurons projecting to the NAc and to the DLS constitute two non-overlapping neuronal populations (**Figure 7E**). Furthermore, we used optogenetics combined with retrograde tracing to investigate which subpopulation of VTA DAT^+^ neurons is mainly functionally connected with the SC. We injected retrograde AAVrg-FLEX-tdTomato in either DLS or NAc in DAT-Cre mice to label distinct VTA DAT^+^ projecting neurons. Within the same animals, we injected AAV-hSyn-ChR2-eYFP in the SC to assess biases in input connectivity based on output-specificity. We performed whole cell patch clamp recordings from identified VTA DAT^+^ projecting neurons (**Figure 7F**) and found that SC makes functional synapses onto VTA DAT^+^ neurons projecting to the DLS with 46.42 % of excitatory connections while VTA DAT^+^-NAc are poorly connected with the SC cells (**Figures 7G, H).** Interestingly, the VTA DAT^+^ neurons connected to SC are located in very lateral position of the VTA as previously shown with the general connectivity of the Figure 1K (**Figure 7I-I’**). Finally, since it has been shown that mPFC makes monosynaptic inputs onto NAc-projecting VTA DAT^+^ neurons (Beier et al. 2015), we hypothesized that VTA DAT^+^ – DLS and VTA DAT^+^ – NAc pathways play different roles during conspecific interaction. Thereby, photostimulation of the VTA DAT^+^ – DLS pathway would mimic the effects presented previously by stimulating SC – VTA. To confirm this hypothesis, we injected Cre-dependent ChR2 (AAV5-EF1α-DIO-ChR2-eYFP) or Cre-dependent eYFP (AAV5-Ef1α-DIO-eYFP) in the VTA of DAT-Cre mice, followed by fiber optic implantation over either DLS or NAc (**Figure 8A-B**). Experimental mice underwent the same behavioural protocol of free social interaction and optogenetic stimulation described previously (**Figure 8C**). As expected, photostimulation of VTA DAT^+^–NAc pathway in ChR2-expressing mice increased time interaction between conspecifics while no changes were observed in eYFP-control group (**Figure 8D-E** and **G**)*(4)*. Remarkably, photostimulation of VTA DAT^+^– DLS in ChR2-expressing mice decreases time sniffing the unfamiliar conspecific (**Figures 8D** and **F-G**), recapitulating the effects of SC – VTA pathway stimulation. Altogether these data indicate that mPFC – VTA^DAT+^ – NAc and SC – VTA^DAT+^ – DLS play distinct but complementary roles in social interaction and orienting response.

**Figure 7:**
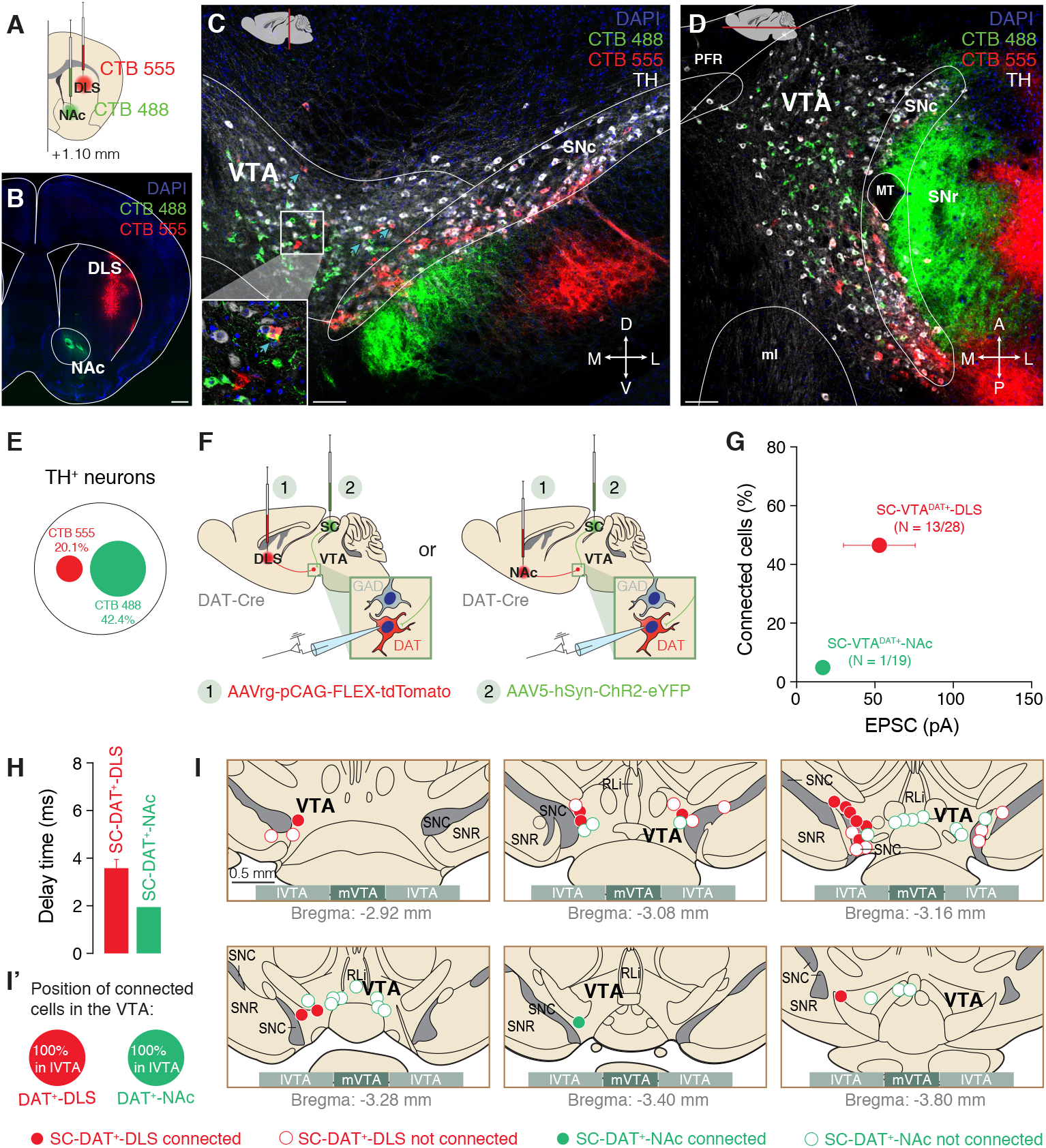
SC is mainly connected to VTA DAT^+^ neurons which project to DLS. **(A)** Schema of injection of CTB 488 and CTB 555 respectively in the Nucleus Accumbens (NAc) or Dorso-Lateral Striatum (DLS). **(B)** Representative image of the NAc and DLS in coronal slice infected with the CTB 488 and CTB 555 (scale bar = 500 *μ*m). **(C)** Coronal plane image of the VTA with TH staining (white) and infected cells projecting to the NAc or DLS (scale bar = 100 *μ*m). **(D)** Horizontal plane image of the VTA with TH staining (white) and infected cells projecting to the NAc or DLS (scale bar = 100 *μ*m). **(E)** Proportion of TH^+^ and either TH^+^/CTB488^+^ (NAc-projecting VTA DAT^+^ neurons) or TH^+^/CTB555^+^ (DLS-projecting VTA DAT^+^ neurons). **(F)** Schema of injections in DAT-Cre mice. The SC was infected with AAV5-hSyn-ChR2-eYFP and the NAc or the DLS with the AAVrg-pCAG-FLEX-tdTomato. Whole cell patch was than performed in DAT^+^ neurons projecting to two different regions. **(G)** Quantification of connected cells from SC onto VTA DAT^+^-NAc or DAT^+^-DLS neurons in relation of the amplitude of EPSC. The VTA DAT neurons receiving projections from the SC are mainly projecting to the DLS with the highest current amplitude. **(H)** Delay time of the EPSC for VTA DAT^+^-NAc or DAT^+^-DLS neurons. **(I-I’)** Position of some patched VTA DAT^+^-NAc or DAT^+^-DLS neurons. SC-VTA^DAT+^-DLS connected neurons are mainly in the lateral part of the VTA (lVTA). N indicates number of cells. Error bars report s.e.m.

**Figure 8:**
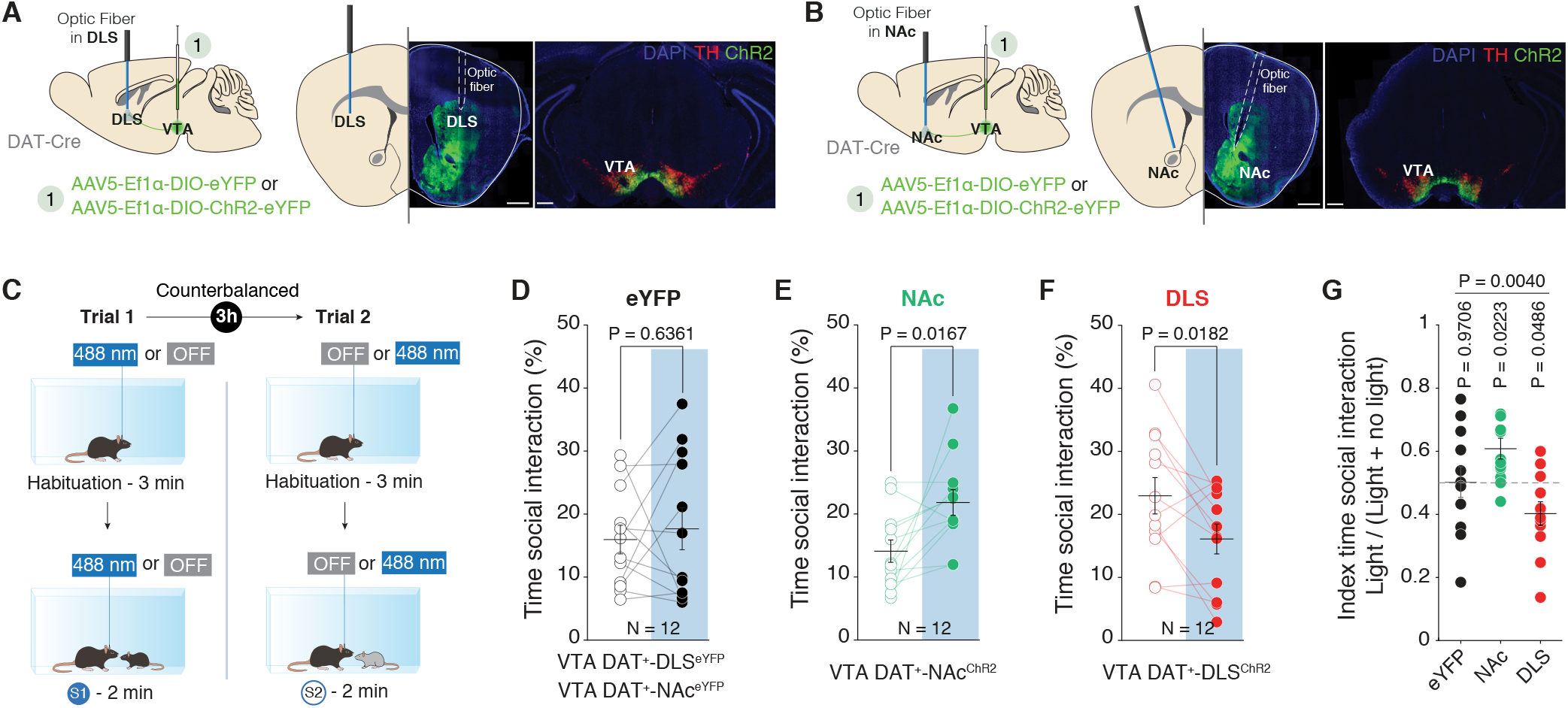
Stimulation of VTA DA – DLS pathway induces similar social interaction alterations observed by stimulating SC – VTA terminals. **(A)** Left: Schema of injections sites in the VTA of DAT-Cre mice with AAV5-Ef1α-DIO-eYFP or AAV5-Ef1α-DIO-ChR2-eYFP, and optic fiber implantation above the dorso-lateral striatum (DLS). Middle: representative image showing the fiber optic’s track in the DLS. Right: representative image of the site of injection in the VTA. (**B**) Left: Schema of injections sites in the VTA of DAT-Cre mice with AAV5-Ef1α-DIO-ChR2-eYFP or AAV5-Ef1α-DIO-eYFP, and optic fiber implantation above the nucleus accumbens (NAc). Middle: representative image showing the fiber optic’s track in the NAc. Right: representative image of the site of injection in the VTA. (**C**) Schema of free social interaction. The mice interact freely with two different unfamiliar mice under both stimulation conditions. **(D)** VTA DAT^+^-NAc^eYFP^ and VTA DAT^+^-DLS^eYFP^ mice do not change time of social interaction between the two stimulation conditions. Paired t test (t_(11)_ = −0.4866). **(E)** VTA DAT -NAc mice increase the + ChR2 time of social interaction during light ON condition. Paired t test (t_(11)_ = −2.8177). **(F)** VTA DAT^+^-DLS^ChR2^ mice decrease the time of social interaction during light ON condition. Paired t test (t_(11)_ = −2.7696). **(G)** Index of time social interaction when photo-stimulating the VTA DAT+ to NAc or DLS pathways. Mice increase social interaction for the NAc pathway while they decrease the social interaction for the DLS pathway. One-way ANOVA (Group main effect: F_(2, 33)_ = 6.56, P = 0.0040) followed by one-sample t test Bonferroni-Holm corrected. N indicates number of cells. Error bars report s.e.m.

## Discussion

The present study examines the role of the SC – VTA pathway during unfamiliar conspecific interaction. We parsed the involvement of two different VTA pathways in distinct, but not mutually exclusive, components of social interaction: while SC – VTA pathway encodes orienting response towards an unfamiliar conspecific, mPFC – VTA circuit activates during investigation of the social stimuli.

It is well established that individual exhibit orienting responses toward novel, unexpected and salient stimuli in the environment *(6)*. Once the salient stimulus is detected, its identity and its potential behavioural relevance need to be considered in order to decide whether to approach or avoid it. In the context of social behaviour, an individual need to detect the presence of other conspecifics and then estimates the valence of a hypothetical interaction, in order to make decisions on whether to initiate or avoid the contacts. After the first interaction, an individual updates the neuronal representation of the value associated with that interaction to guide its subsequent decisions *(24)* to or -not-to maintain that social contact. Although it has been previously suggested that deficits in orienting response may underlie part of the deficits in social interaction associated with Autism Spectrum Disorders *(25)* the importance of orienting response in social interaction has been largely ignored. Here we showed for the first time that VTA not only play an important role during social interaction *(26)*, but that different inputs onto this region convey different signals. Among the inputs, we showed that SC – VTA pathway encodes orienting response toward salient stimuli, and alterations of its activity affects social interaction.

SC is an evolutionary ancient structure organized in functionally and anatomically distinct layers. While the upper layers are exclusively visual, the medial and deeper layers have multisensory and motor functions *(27)*. It has been suggested that this structure, *via* different output projections, mediates both orienting and avoidance responses to novel sensory stimuli *(15, 17)*. Moreover, SC lesions generally result in deficits in visual orientation, sensory neglect *(28)* and aberrant predatory responses *(29)*. Previous reports have shown that SC mediates visually-induced defensive behaviours. Indeed, the activity of excitatory neurons in the deep layers of the medial SC represents the saliency of the threat stimulus and is predictive of escape behaviour *(30, 31)*. More specifically, the SC projections onto the VTA GABA neurons would promote “flight” behavior in threatening context *(19)*.

Here, we demonstrated, for the first time, the importance of the SC in social contexts. Indeed, our data show that SC – VTA pathway is transiently activated during orientation toward an unfamiliar conspecific. Perturbing the physiological activity pattern of this pathway affects orientation and, consequently, social interaction. Thus, the activity of SC – VTA pathway is tightly tuned to environmental stimuli to heighten attention and favour orienting response.

Midbrain DA neurons are located in two neighbouring nuclei: the Substantia Nigra (SN) and the VTA *(32)* from where they project diffusely throughout the brain. Canonically, DA neurons are involved in reward-based learning, yet some studies observed that a subpopulation is activated by aversive stimuli *(33, 34, 35)*. These findings have led to the hypothesis that these DA neurons may signal “incentive salience” and facilitate a behavioural response when a salient stimulus is detected *(36, 23)*. Interestingly, it has been shown that short latency phasic responses can be elicited in DA neurons by unexpected rewards or by conditioned stimuli that predict the reward *(37, 36, 38)*. In this regard, different studies have shown that the midbrain SC directly projects to midbrain DA neurons and can relay reward-predictive sensory information to them *(13, 39)*. In our study, we prove that SC projects to a subpopulation of VTA DA neurons that sends outputs to the DLS, thus pointing at the DA neurons as a relay node between the SC and the striatum. Interestingly, the looped circuit connecting the SC to the basal ganglia may be a “candidate mechanisms to perform the pre-attentive selections required to determine whether gaze should be shifted, and if so, to which stimulus”*(18)*. Our data therefore propose that, in the context of unfamiliar conspecific interaction, signals from the SC to the VTA DA neurons trigger orientation towards salient stimuli to favour interaction.

Social behaviour involves a sequence of different actions: from interruption of the ongoing behaviour, stimulus evaluation, decision-making and further exploration. In particular, social motivation is characterized by preferential orientation towards social stimuli, evaluation of the rewarding properties of conspecific interaction and effort to maintain social bonds *(26)*. Social motivation deficits play a central role in Autism Spectrum Disorders (ASDs), a neuropathology characterized by major social behaviour alterations. Although evidence suggests that VTA is central to social deficits in ASD patients *(40)*, the origins of these deficits are still largely unknown. Interestingly, one of the core diagnostic criteria for ASD includes decrease eye-contact *(41)* and eye-tracking experiments have shown an impairment in orientation towards social stimuli in ASD patients *(42)*. Here, our data support the hypothesis that a precise activation of inputs to the VTA is necessary to promote social orientation and then interaction. Indeed, by using ChR2 within the SC – VTA pathway, we disrupt the time lock increase in activity during the orientation toward a conspecific. This manipulation *per se* is sufficient to result in altered social interaction. We therefore propose that intact functionality of the SC – VTA^DA^ – DLS pathway may be fundamental for orientation towards conspecifics and deficits in this pathway may be upstream to social motivation dysfunctions in ASD patients.

In conclusion our data strongly suggest while SC – VTA^DA^ – DLS pathway encodes orienting response towards an unfamiliar conspecific, mPFC – VTA^DA^ – NAc circuit function regulates the investigation and maintenance of social interaction. Elucidating the brain circuits underlying conspecific interaction is therefore essential, not only to understand how social behaviour occurs, but also to comprehend the aberrant neural mechanisms underlying social deficits in psychiatric disorders.

## Material and Methods

### Mice

Male wild type (WT; C57Bl/6J), DAT-iresCre (Slc6a3^tm1.1(cre)Bkmn^/J, called DAT-Cre in the rest of manuscript) and GAD2-iresCre (Gad2^tm2(creZjh)^/J, called GAD65-Cre in the rest of manuscript) were employed for this study. Mice were housed in groups (weaning at P21 – P23) under a 12 hours light – dark cycle (7:00 a.m.–7:00 p.m.) with food and water *ad libidum*. All physiology and behaviour experiments were performed during the light cycle (the experiments were performed in a time window that started approximately 2 h after the end of the dark circle and ended 2 h before the start of the next dark circle) and were conducted in a room with fixed illumination (20 Lux), temperature (22-24°C) and humidity (40%). All the procedures performed at UNIGE complied with the Swiss National Institutional Guidelines on Animal Experimentation and were approved by the respective Swiss Cantonal Veterinary Office Committees for Animal Experimentation.

### Stereotaxic injection and optic fiber implantation

#### Optogenetic experiments

rAAV5-hSyn-hChR2(H134R)-eYFP, rAAV5-hSyn-Jaws-KGC-GFP-ER2 or rAAV5-hSyn-eYFP were injected in WT mice at 7-8 weeks. Mice were anesthetized with a mixture of oxygen (1 L/min) and isoflurane 3% (Baxter AG, Vienna, Austria). The skin was shaved, locally anesthetized with 40–50 *μ*L lidocaine 0.5% and disinfected. The animals were placed in a stereotactic frame (Angle One; Leica, Germany) and bilateral injections were performed in the SC (ML ± 0.8 mm, AP −3.4 mm, DV −1.5 mm from Bregma, 500 nL per side). The virus was injected via a glass micropipette (Drummond Scientific Company, Broomall, PA). The virus was incubated for 3 – 4 weeks and subsequently mice were implanted with optic fibers above the VTA. The animals were anesthetized, placed in a stereotactic frame, the skin was shaved and a unilateral craniotomy was performed above the VTA. The optic fiber was implanted with a 10° angle at the following coordinates: ML ± 0.9 mm, AP −3.2 mm, DV −3.95 mm from Bregma above the VTA and fixed to the skull with dental acrylic. For optogenetic experiments using DAT-Cre mice, the animals were injected using the protocol described above with rAAV5-Ef1α-DIO-hChR2(H134R)-eYFP or rAAV5-Ef1α-DIO-eYFP in the VTA (ML ± 0.5 mm, AP −3.2 mm, DV −4.25/-4.00 mm from Bregma). The virus was incubated for 3 – 4 weeks and subsequently mice were implanted with optic fibers above either the Nucleus Accumbens (NAc) with 15° angle at the following coordinates: ML ± 2.0 mm, AP +1.2 mm, DV −4.2 mm from Bregma; or the dorsolateral Striatum (DLS) at the following coordinates: ML ± 2.0 mm, AP +1.0 mm, DV −2.5 mm from Bregma. Injections and implantation sites were confirmed post hoc.

#### Fiber photometry experiments

AAVrg-Ef1α-mCherry-IRES-Cre injections in the VTA (500 nL in total) and unilateral AAV9-hSyn-FLEX-GCaMP6s-WPRE-SV40 injection unilaterally in the SC (500 nL: ML ± 0.8 mm, AP –3.4 mm, DV −1.5 mm from Bregma) or mPFC (500 nL: ML ± 0.3 mm, AP + 1.95 mm, DV −2.1 mm from Bregma) in adult WT mice were conducted following the protocol described above. The viruses were incubated 3 – 4 weeks prior optic fibers implantation. The optic fibers were unilaterally implanted following the above-mentioned protocol in the SC (ML ± 0.8 mm, AP –3.4 mm, DV −1.5 mm from Bregma) and mPFC (ML ± 0.3 mm, AP + 1.95 mm, DV −2.1 mm from Bregma). Injections and implantation sites were confirmed post hoc.

#### E*x-vivo* electrophysiological recording experiments

Injections of rAAV5-hSyn-DIO-mCherry, AAVrg-pCAG-FLEX-tdTomato-WPRE, rAAV5-hSyn-hChR2(H134R)-eYFP, rAAV5-hSyn-hChR2(H134R)-mCherry and/or rAAV5-hSyn-Jaws-KGC-GFP-ER2, were performed in WT, DAT-Cre or GAD65-Cre mice (depending on the experiment). rAAV5-hSyn-hChR2(H134R)-eYFP and rAAV5-hSyn-hChR2(H134R)-mCherry + rAAV5-hSyn-Jaws-KGC-GFP-ER2 were injected in the SC, at the same coordinates previously described. rAAV5-hSyn-DIO-mCherry was injected in the VTA at the same coordinates previously described. AAVrg-pCAG-FLEX-TdTomato was bilateraly injected either in the NAc at these coordinates: ML ± 1.0 mm, AP +1.2 mm, DV −4.4 / −4.0 mm from Bregma, 500 nL each side; either the DLS at following coordinates: ML ± 2.0 mm, AP +1.0 mm, DV −2.8 mm from Bregma, 500 nL each side. The viruses were incubated 3 – 4 weeks before to perform *ex-vivo* electrophysiological recordings.

#### Anatomical validation experiments

Bilateral injections of rAAV5-hSyn-eYFP (500 nL per side) in the SC were performed as described above. CAV2-Cre injection in the VTA (500 nL per side) and rAAV5-hSyn-DIO-mCherry injection in the SC (500 nL per side) were conducted following the protocol described above at the already mentioned coordinates. WT mice were bilaterally injected using Cholera Toxin subunit-B Alexa fluor 555 (CTB 555) or CTB 488 respectively in the DLS (ML ± 2.0 mm, AP +1.0 mm, DV −2.8 mm from Bregma, 200 nL each side) and the NAc (ML ± 1.0 mm, AP +1.2 mm, DV −4.4 / −4.0 mm from Bregma, 200 nL each side). The CTB 488 and CTB 555 were incubated during 2 weeks before immunostaining procedures.

### Free interaction during *in-vivo* calcium imaging

An arena similar to the animal’s homecage was used for the free social interaction task. The arena was cleaned using 70% ethanol and the bedding replaced after each trial. Mice injected with AAVrg-Ef1α-mCherry-IRES-Cre in the VTA and AAV9-hSyn-FLEX-GCaMP6s-WPRE-SV40 in the SC or mPFC were first placed in the arena and were free to explore the new environment for 5 mins. After this habituation period, a social stimulus (unfamiliar juvenile conspecific sex-matched C57BL/6J, 3 – 4 weeks) was introduced in the cage, and the animals were free to interact during 5 mins. During the entire session the bulk calcium waves of SC – or mPFC – VTA projecting neurons was recorded with 1-site fiber photometry system (Doric lenses).

Every session was video-tracked and recorded using Ethovision XT (Noldus, Wageningen, the Netherlands). Non-aggressive interaction was manually scored. After the task, the animals were sacrificed and the viral infection was verified.

### Free interaction during optogenetic manipulation

Similarly to the protocol described previously, an arena similar to the animal’s homecage was used for the free social interaction task. The arena was cleaned using 70% ethanol and the bedding replaced after each trial. All SC eYFP-, ChR2 and Jaws-expressing mice underwent both conditions of light-ON or light-OFF epochs. The light stimulation or inhibition protocols were applied depending on a random assignment and lasted during all the trial. For eYFP-control and ChR2-expressing mice, the following optogenetic stimulation protocol was used: burst of 8 pulses of 4 msec light at 30 Hz every 5 sec (Imetronic, Pessac, France), wavelength of 488 nm (BioRay Laser, Coherent). For eYFP-control and Jaws-expressing mice, a constant light-ON epoch was delivered (Imetronic, Pessac, France), at 640 nm wavelength (BioRay Laser, Coherent). The experimental mice were first placed in the arena and were free to explore the new environment for 3 mins. After this period, a social stimulus (unfamiliar juvenile conspecific sex-matched C57BL/6J, 3 – 4 weeks; S1) was introduced in the cage, and the animals were free to interact during 2 mins. The mice were then removed and placed in their respective cages. After 3 hours, the same experimental design was performed and the mice that received the light stimulation or inhibition protocols did not receive it and *vice versa*. Another unfamiliar conspecific stimulus (S2) was introduced for 2 mins in the cage after the 3 mins of exploration. The power expected at the tip of the optic fiber was between 8 – 12 mW. To this aim the laser power was checked before every experiment and the power at the optic fiber tip was controlled before any implantation.

Likewise, DAT-Cre mice injected with rAAV5-Ef1α-DIO-eYFP or rAAV5-Ef1α-DIO-hChR2(H134R)-eYFP in the VTA and implanted with double optic fiber either in the Nucleus Accumbens (VTA DAT^+^-NAc^eYFP^ or VTA DAT^+^-Nac^ChR2^) or Dorsolateral Striatum (VTA DAT^+^-DLS^eYFP^ or VTA DAT^+^-DLS^ChR2^) underwent the same protocol of free social interaction task as described above.

Every session was video-tracked and recorded using Ethovision XT (Noldus, Wageningen, the Netherlands). Non-aggressive interaction was manually scored (experimenter blind to the viral injection) when the experimental mouse initiated the action and when the nose of the animal was touching the juvenile conspecific. Using the vector formed by the gravity center and the nose of the experimental mice, it has been possible to calculate the head orientation towards the gravity center of the conspecific stimulus.

After the task, the animals were sacrificed and the viral infection was verified. If no infection was detected or the optic fiber not in the right area, the mice were excluded from the batch.

### Social and non-social orientation task during *in-vivo* calcium imaging

The arena of the orientation task is composed of two cylinders (height = 25 cm) positioned one inside the other. The smaller cylinder (Ø = 8 cm) is composed by transparent plastic and present small holes (Ø = 0.3 cm) which prevent social contacts but allow olfactory, auditory and visual cues. The mice injected with AAVrg-Ef1α-mCherry-IRES-Cre in the VTA and AAV9-hSyn-FLEX-GCaMP6s-WPRE-SV40 in the SC or mPFC were gently placed in the small cylinder which allow the experimental animal only to turn left or right. After a 5-mins period of habituation, a social stimulus (unfamiliar juvenile conspecific sex-matched C57BL/6J, 3 – 4 weeks) or a moving ball (64.60, Sphero Mini, Sphero Edu) was introduced between the small and the big cylinder (Ø = 20 cm). The social stimulus or the moving ball were allowed to freely move in the inter-cylinder space for 5 minutes. The ball was programmed to move similarly to a conspecific, with approximately the same speed. During the entire session the bulk calcium waves of SC- or PFC -VTA projecting neurons was recorded with 1-site fiber photometry system (Doric lenses). Every session was video-tracked and recorded using Ethovision XT (Noldus, Wageningen, the Netherlands). Ipsi- and contra-recorded orientation and passive crossing were manually scored. The arena was cleaned using 70% ethanol between each trial. After the task, the animals were sacrificed and the viral infection was verified.

### Orientation task during optogenetic manipulation

The arena described previously was used during optogenetic experiments. All SC eYFP-, ChR2 and Jaws-expressing mice underwent both conditions of light-ON or light-OFF epochs. The light stimulation or inhibition protocols were applied depending on a random assignment and lasted during all the trial. For eYFP-control and ChR2-expressing mice, the following optogenetic stimulation protocol was used: burst of 8 pulses of 4 msec light at 30 Hz every 5 sec (Imetronic, Pessac, France), wavelength of 488 nm (BioRay Laser, Coherent). For eYFP-control and Jaws-expressing mice, a constant light-ON epoch was delivered (Imetronic, Pessac, France), at 640 nm wavelength (BioRay Laser, Coherent). The experimental mice were gently placed in the small cylinder allowing the experimental animal only to turn left or right. After a 5-mins period of habituation, a social stimulus (unfamiliar juvenile conspecific sex-matched C57BL/6J, 3 – 4 weeks, S1) was introduced between the small and the big cylinder. The social stimulus was allowed to freely move in the inter-cylinder space for 2 minutes. The mice were then removed and placed in their respective cages. After 3 hours, the same experimental design was performed and the mice that received the light stimulation or inhibition protocols did not receive it and *vice versa*. Another unfamiliar conspecific stimulus (S2) was introduced for 2 mins in the inter-cylinder space after the 5-mins period of habituation. The power expected at the tip of the optic fiber was between 8 – 12 mW. To this aim the laser power was checked before every experiment and the power at the optic fiber tip was controlled before any implantation. The arena was cleaned using 70% ethanol between each trial.

Every session was video-tracked and recorded using Ethovision XT (Noldus, Wageningen, the Netherlands). Using the vector formed by the gravity center and the nose of the experimental mice, it has been possible to calculate the head orientation towards the gravity center of the conspecific stimulus and the relative position of the social stimulus.

After the task, the animals were sacrificed and the viral infection was verified. If no infection was detected or the optic fiber not in the right area, the mice were excluded from the batch.

### Analyses of fiber photometry data

Fiber photometry data were analyzed with custom MatLab codes. Raw signals were recorded and adjusted according to the overall trend to take account the photo-bleaching. For each experiment we defined the F_0_ as the 5 mins baseline activity before presentation of the social stimulus. The fluorescence change was determined as ΔF/F (Z-score) and calculated as (F-F_0_)/F_0_ where F is the fluorescence at each bin. The acquisition frequency was at 12kHz (bins of 1/12158 sec) for the entire recordings. The construction of Peri-event time histogram (PETH) was made by aligning and centering specific events. These events were obtained by manual scoring of specific behaviours at specific times to link Ca^2+^ activity with the events. After normalization of ΔF/F, a convolution using a Kernel-Gaussian sliding window of 2000 bins was applied on the data (*gausswin* MatLab function).

### Open Field

Mice injected with rAAV5-hSyn-eYFP, rAAV5-hSyn-hChR2(H134R)-eYFP or rAAV5-hSyn-Jaws-KGC-GFP-ER2 in SC and implanted with optic fiber above the VTA performed the open-field task (OF). The mice were placed in the OF arena for 10 min. The apparatus consisted in a 40 cm sided Plexiglas squared arena. The mice were randomly assigned to the optical light stimulation/ inhibition protocols (as previously described)(Imetronic system, Pessac, France). After 10 min of free exploration the mice were placed back into their homecage. 3 hours later the animals were re-tested in the OF experiment and the mice that received the light-ON epoch did not receive it and *vice versa*. At the end, all the animals performed both conditions. The OF task was video-tracked (Ethovision, Noldus, Wageningen, the Netherlands) to automatically obtain the distance moved and the time passed in the center. After the task, the animals were sacrificed and the viral infection was verified. If no infection was noticed or the optic fiber not in the right area, the mice were excluded from the batch. The apparatus was cleaned using 70% ethanol after each session.

### Real Time Place Preference task

WT mice injected with rAAV5-hSyn-eYFP, rAAV5-hSyn-hChR2(H134R)-eYFP or rAAV5-hSyn-Jaws-KGC-GFP-ER2 in SC and implanted with optic fiber in the VTA performed the real time place preference (rtPP). The rtPP experiment was conducted in an apparatus (spatial place preference; BioSEB) consisting of two adjacent chambers (20 × 20 × 25 cm) with dot (black) or stripe (grey) wall patterns, connected by a lateral corridor (7 × 20 × 25 cm) with transparent walls and floor. The dot chamber was always associated to rough floor, while the stripe chamber with smooth floor. The illumination level was uniform between the two chambers and set at 10 – 13 lux. Imetronic (Pessac, France) tracking software was used to track animal’s movements, the time spent within each chamber and to deliver optogenetic protocols. The mice were placed in the apparatus and were free to explore both chambers during 10 min. One chamber was systematically associated with a high bursting optogenetic stimulation protocol for ChR2-expressing mice (burst of 5 pulses of 4 msec at 20 Hz every 250 msec) while the other was not associated with any optical stimulation protocol. The chamber associated with the light stimulation or inhibition was counterbalanced to avoid any internal bias due to the cues of the compartment. The OF task was video-tracked (Ethovision, Noldus, Wageningen, the Netherlands) to automatically obtain the time passed in each chamber of the mice.

After the task, the animals were sacrificed and the viral infection was verified. If no infection was noticed or the optic fiber not in the right area, the mice were excluded from the batch. The apparatus was cleaned using 70% ethanol after each session.

### Ex vivo slice physiology

200 – 250 *μ*M thick horizontal midbrain slices were prepared from C57Bl/6J WT, GAD65-cre or DAT-Cre mice. Brains were sliced by using a cutting solution containing: 90.89 mM choline chloride, 24.98 mM glucose, 25 mM NaHCO_3_, 6.98 mM MgCl_2_, 11.85 mM ascorbic acid, 3.09 mM sodium pyruvate, 2.49 mM KCl, 1.25 mM NaH_2_PO_4_, and 0.50 mM CaCl_2_. Brain slices were incubated in cutting solution for 20 – 30 min at 35°. Subsequently, slices were transferred in artificial cerebrospinal fluid (aCSF) containing: 119 mM NaCl, 2.5 mM KCl, 1.3 mM MgCl_2_, 2.5 mM CaCl_2_, 1.0 mM NaH_2_PO_4_, 26.2 mM NaHCO_3_, and 11 mM glucose, bubbled with 95% O_2_ and 5% CO_2_) at room temperature. Whole-cell voltage clamp or current clamp electrophysiological recordings were conducted at 32° – 34° in aCSF (2 – 3 ml.min^−1^, submerged slices). Patch pipettes were filled with a Cs^+^-based low Cl^−^ internal solution containing 135 mM CsMeSO_3_, 10 mM HEPES, 1 mM EGTA, 3.3 mM QX-314, 4 mM Mg-ATP, 0.3 mM Na_2_-GTP, 8 mM Na_2_-Phosphocreatine (pH 7.3 adjusted with CsOH; 295 mOsm). For ChR2-expressing WT mice, SC neurons were patched and the light stimulation protocol was assessed. For DAT-Cre and GAD65-Cre mice DAT^+^ or GAD^+^ neurons respectively of the VTA were identified as mCherry^+^ cells. Brief pulses of blue light (10 msec) were delivered at the recording site at 10 sec intervals under control of the acquisition software. Cells were helded at −60mV in order to evoke excitatory post-synaptic currents (EPSCs) or at 0 mV to evoke inhibitory post-synaptic currents (IPSCs). The same protocol was in DAT-Cre mice injected with the AAVrg-pCAG-FLEX-TdTomato on the NAc or DLS, and the DAT^+^ were identified as TdTomato^+^ cells.

To confirm the terminal inhibition of the AAV5-hSyn-Jaws-KGC-GFP-ER2 onto VTA DA neurons from SC infected cells, we injected a mix of AAV5-hSyn-Jaws-KGC-GFP-ER2 and AAV5-hSyn-hChR2(H134R)-mCherry in the SC and the AAV5-hSyn-DIO-mCherry in the VTA of DAT-Cre mice. Brief pulses of blue light (10 msec) were delivered at 10 sec intervals during 2 mins then brief pulses of red light (30 msec) and blue light (10 msec) were applied at the same time for 10 mins.

### Immunohistochemistry and cell counting

Infected mice were anesthetized with pentobarbital (Streuli Pharma) and sacrificed by intra-cardial perfusion of 0.9% saline followed by 4% paraformaldehyde (PFA; Biochemica). Brains were post-fixed overnight in 4% PFA at 4°C. 24 hours later, they were washed with PBS and then 50 μm thick sliced with a vibratome (Leica VT1200S).

Previously prepared slices were washed three times with phosphate buffered saline (PBS) 0.1M. Brain slices were pre-incubated with PBS-BSA-TX buffer (10% BSA, 0.3% Triton X-100, 0.1% NaN_3_) for 60 min at room temperature in the dark. Subsequently, cells were incubated with primary antibodies diluted in PBS-BSA-TX (3% BSA, 0.3% Triton X-100, 0.1% NaN_3_) overnight at 4°C in the dark. The following day cells were washed three times with PBS 0.1M and incubated for 60 min at room temperature in the dark with the secondary antibodies diluted in PBS-Tween buffer (0.25% Tween-20). Finally, slices were mounted using Fluoroshield mounting medium with DAPI (abcam, Cat#ab104139). In this study, the following primary antibodies were used: mouse monoclonal anti-CaMKII alpha (1/100 dilution, ThermoFisher, Cat#MA1-048, RRID: AB_325403), rabbit polyclonal anti-mCherry (1/200 dilution, abcam, Cat#ab167453) and rabbit polyclonal anti-Tyrosine Hydroxylase (1/500 dilution, abcam, Cat#ab6211). The following secondary antibodies were used at 1/500 dilution: donkey anti-mouse Alexa Fluor 488 (ThermoFisher, Cat#R37114, RRID: AB_2556542), donkey anti-rabbit Alexa Fluor 555 (ThermoFisher, Cat#A32794, RRID: AB_2762834) and donkey anti-rabbit Alexa Fluor 647 (ThermoFisher, Cat# A32795, RRID: AB_2762835). Immunostained slices were imaged using the confocal laser scanning microscopes Zeiss LSM700 and LSM800. Larger scale images were taken with the widefield Axioscan.Z1 scanner.

Cell counting of mCherry^+^ cells was performed on 50μm thick SC slices from 3 SC^AAV-DIO-mCherry^-VTA^CAV-Cre^ mice (5 slices for each animal). For each slice, images from the SC and PAG were acquired bilaterally along the whole SC dorso-ventral axis. The mCherry^+^ cells were counted automatically using MetaMorph Software (Molecular Devices). The total percentage of cells located in the different layers of the SC or the PAG was calculated by averaging the total number mCherry^+^ of each mouse. After CaMKIIa immunostaining onto SC slices, the number of mCherry^+^/ CaMKIIa^+^ and mCherry^+^/ CaMKIIa^−^ cells were counted.

### Viruses

rAAV5-hSyn-hChR2(H134R)-eYFP (Titer ≥ 7×10^12^ vg.mL^−1^, Addgene), rAAV5-hSyn-Jaws-KGC-GFP-ER2 (Titer ≥ 3.8×10^12^ vg.mL^−1^, UNC Vector Core), rAAV5-hSyn-eYFP (Titer ≥ 7×10^12^ vg.mL^−1^, Addgene), rAAV5-hSyn-hChR2(H134R)-mCherry (Titer ≥ 7×10^12^ vg.mL^−1^, Addgene), rAAV5-Ef1α-DIO-hChR2(H134R)-eYFP (Titer ≥ 4.2×10^12^ vg.mL^−1^, UNC Vector Core), rAAV5-Ef1α-DIO-eYFP (Titer ≥ 4.2×10^12^ vg.mL^−1^, UNC Vector Core), rAAV5-hSyn-DIO-mCherry (Titer ≥ 7×10^12^ vg.mL^−1^, Addgene), AAVrg-pCAG-FLEX-tdTomato-WPRE (Titer ≥ 1×10^13^ vg.mL^−1^, Addgene), CAV-2 Cre (Titer ≥ 2.5×10^11^ pp, Plateforme de Vectorologie de Montpellier, PVM), AAV9-hSyn-FLEX-GCamp6s-WPRE-SV40 (Titer ≥ 3.8×10^12^ vg.mL^−1^, UNC Vector Core), AAVrg-Ef1α-mCherry-IRES-Cre (Titer ≥ 7×10^12^ vg.mL^−1^, Addgene). Cholera Toxin Subunit B (Recombinant), Alexa Fluor™ 488 Conjugate (ThermoFisher Scientific, C22841), Cholera Toxin Subunit B (Recombinant), Alexa Fluor™ 555 Conjugate (ThermoFisher Scientific, C34776).

### Statistical analysis

No statistical methods were used to predetermine the number of animals and cells, but suitable sample sizes were estimated based on previous experience and are similar to those generally employed in the field. The animals were randomly assigned to each group at the moment of viral infections or behavioural tests. Statistical analysis was conducted with MatLab (The Mathwork), R or GraphPad Prism 7 (San Diego, CA, USA). Statistical outliers were identified by using the criterion *Mean*_*Value*_ ± 3 × *Std*_*Value*_ and excluded from the analysis. The normality of sample distributions was assessed with the Shapiro–Wilk criterion and when violated non-parametrical tests were used. When normally distributed, the data were analysed with independent t test, paired t test, while for multiple comparisons one-way ANOVA and repeated measures (RM) ANOVA were used. When normality was violated, the data were analysed with Mann–Whitney test. For the analysis of variance with two factors (two-way ANOVA, RM two-way ANOVA and RM two-way ANOVA by both factors), normality of sample distribution was assumed, and followed by Bonferroni-Holm correction test or Bonferroni post-hoc test. All the statistical tests adopted were two-sided. When comparing two samples distributions similarity of variances was assumed, therefore no corrections were adopted. Data are represented as the *Mean* ± *s*. *e*. *m*. and the significance was set at P < 0.05.

## Acknowledgments

We are grateful to Lorena Jourdain for the technical support. We thank Manuel Mameli for the constructive comment on the manuscript and the entire Bellone lab for discussion.

## Author Contribution

CB, CPS and AC designed the study. CPS and AC performed the optogenetic experiments, virus injections and optic fiber implantation. AC performed the immunohistochemistry and the fiber photometry experiments and analysed the data with the help of CPS, CH and AC. PE, SB, and SM performed electrophysiological recordings. CB wrote the manuscript with the help of CPS and AC. AC prepared the figures.

## Data and material availability

All the data are in the manuscript or in supplementary material. Videos, behavioural scoring and code analysis will be made available on request.

## Supplementary figures legends

**Supplementary figure 1:**
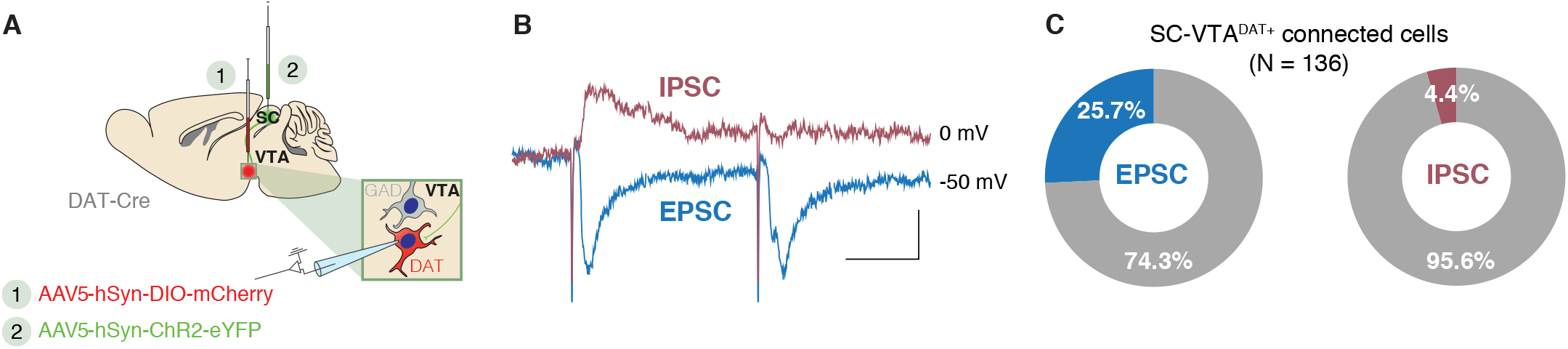
SC prevalently evokes more EPSC than IPSC in VTA DAT^+^ cells. **(A)** Schema of injection in the SC with AAV5-hSyn-ChR2-eYFP and in the VTA with AAV5-hSyn-DIO-mCherry, then patch of the VTA DAT^+^. **(B)** Example traces of optogenetically-induced excitatory post-synaptic current (EPSC) or optogenetically-induced inhibitory post-synaptic current (IPSC) in VTA DAT^+^ neurons. **(C)** Percentage of evoked EPSC or IPSC in the VTA DAT^+^ neurons connected with the SC.

**Supplementary figure 2:**
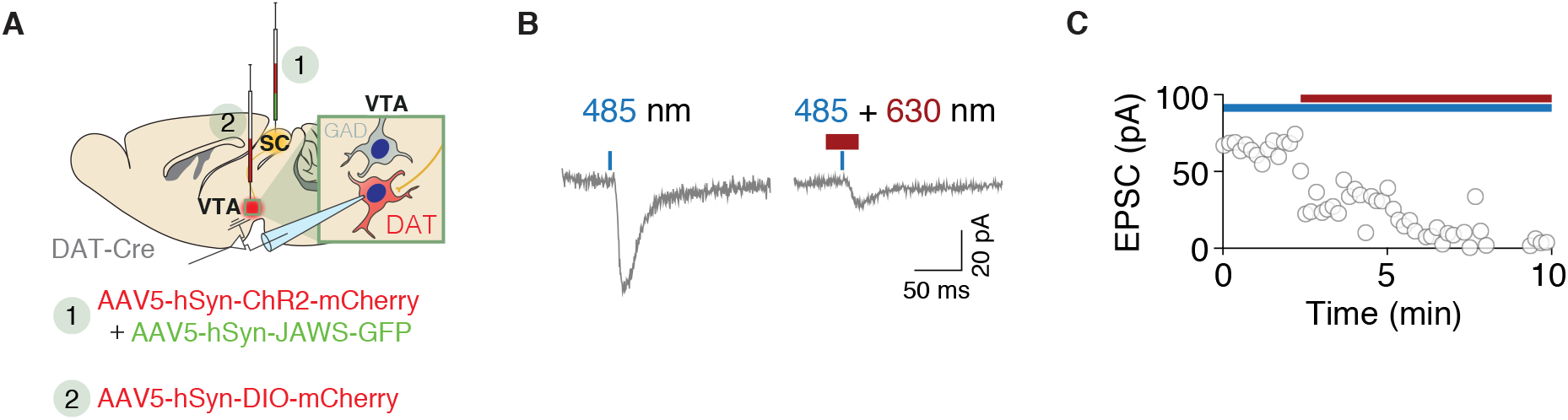
SC Jaws-expressing neurons induce terminal inhibition in VTA DA cells. **(A**) Schema of injections in the SC with AAV5-hSyn-ChR2-mCherry + AAV5-hSyn-JAWS-GFP and with AAV5-hSyn-DIO-mCherry in the VTA. **(B)** Example traces of evoked EPSC after photostimulation (left) and photostimulation followed by photoinhibition (right). **(C)** Amplitude of evoked EPSC in function of the time. Photostimulation is indicated in blue and photoinhibition in red. The graph shows an induced EPSC in the VTA DAT^+^ neurons when the blue light only is shined. However, with contingent shining of blue and red lights, the current approach to 0 confirming a terminal inhibition from the SC onto VTA DAT^+^ neurons.

**Supplementary figure 3:**
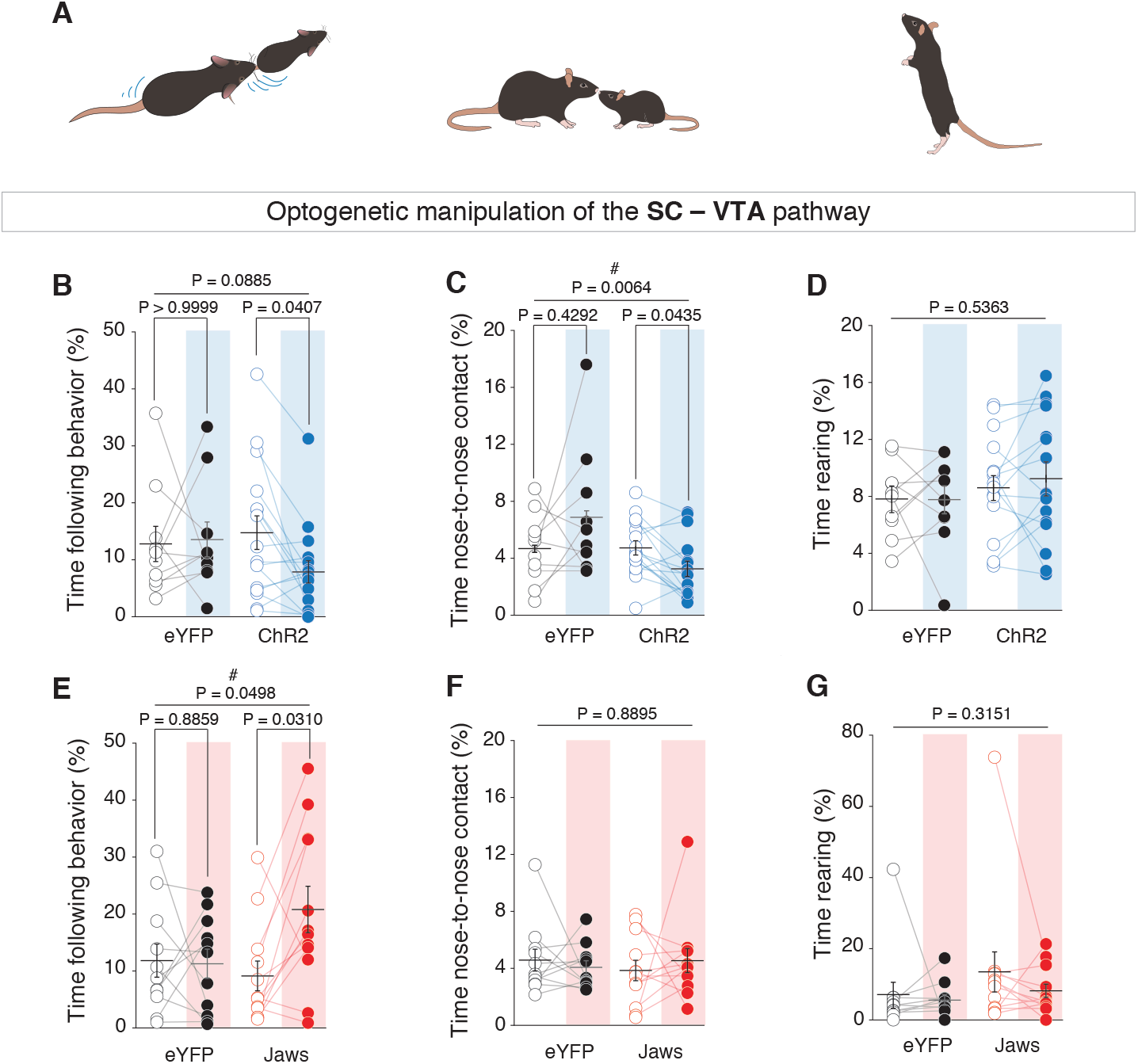
Quantification of nose-to-nose, nose-to-body and rearing events during free social interaction task. **(A)** Schema of a mouse performing nose-to-nose (left), nose-to-body (center) or rearing (right). **(B)** Time of following behavior for SC eYFP and ChR2-expressing mice under light-ON and light-OFF epochs. RM two-way ANOVA (Light main effect: F_(1,24)_ = 3.1522, P = 0.0885; Virus main effect: F_(1,24)_ = 0.3289, P = 0.5716; Interaction Light x Virus: F_(1,24)_ = 2.7778, P = 0.1086) followed by Bonferroni-Holm post-hoc test correction. **(C)** Time of nose-to-nose contact for SC eYFP and ChR2-expressing mice under light-ON and light-OFF epochs. RM two-way ANOVA (Light main effect: F_(1,24)_ = 0.0064, P = 0.9370; Virus main effect: F_(1,24)_ = 4.3794, P = 0.0471; Interaction Light x Virus: F_(1,24)_ = 6.4369, P = 0.0181) followed by Bonferroni-Holm post-hoc test correction. **(D)** Time of rearing behavior during free social interaction task for SC eYFP and ChR2-expressing mice under light-ON and light-OFF epochs. RM two-way ANOVA (Light main effect: F_(1,28)_ = 0.3921, P = 0.5363; Virus main effect: F_(1,28)_ = 1.3238, P = 0.2596; Interaction Light x Virus: F_(1,28)_ = 0.5985, P = 0.4456). **(E)** Time of following behavior for SC eYFP and Jaws-expressing mice under light-ON and light-OFF epochs. RM two-way ANOVA (Light main effect: F_(1,21)_ = 4.3333, P = 0.0498; Virus main effect: F_(1,21)_ = 0.9925, P = 0.3305; Interaction Light x Virus: F_(1,21)_ = 4.7701, P = 0.0404) followed by Bonferroni-Holm post-hoc test correction. **(F)** Time of nose-to-nose contact for SC eYFP and Jaws-expressing mice under light-ON and light-OFF epochs. RM two-way ANOVA (Light main effect: F_(1,21)_ = 0.0198, P = 0.8895; Virus main effect: F_(1,21)_ = 0.0476, P = 0.8294; Interaction Light x Virus: F_(1,21)_ = 0.5243, P = 0.4770). **(G)** Time of rearing behavior during free social interaction task for SC eYFP and Jaws-expressing mice under light-ON and light-OFF epochs. RM two-way ANOVA (Light main effect: F_(1,21)_ = 1.0594, P = 0.3151; Virus main effect: F_(1,21)_ = 1.3808, P = 0.2531; Interaction Light x Virus: F_(1,21)_ = 0.3329, P = 0.5701). # Indicates significant interaction. Error bars report s.e.m.

**Supplementary figure 4:**
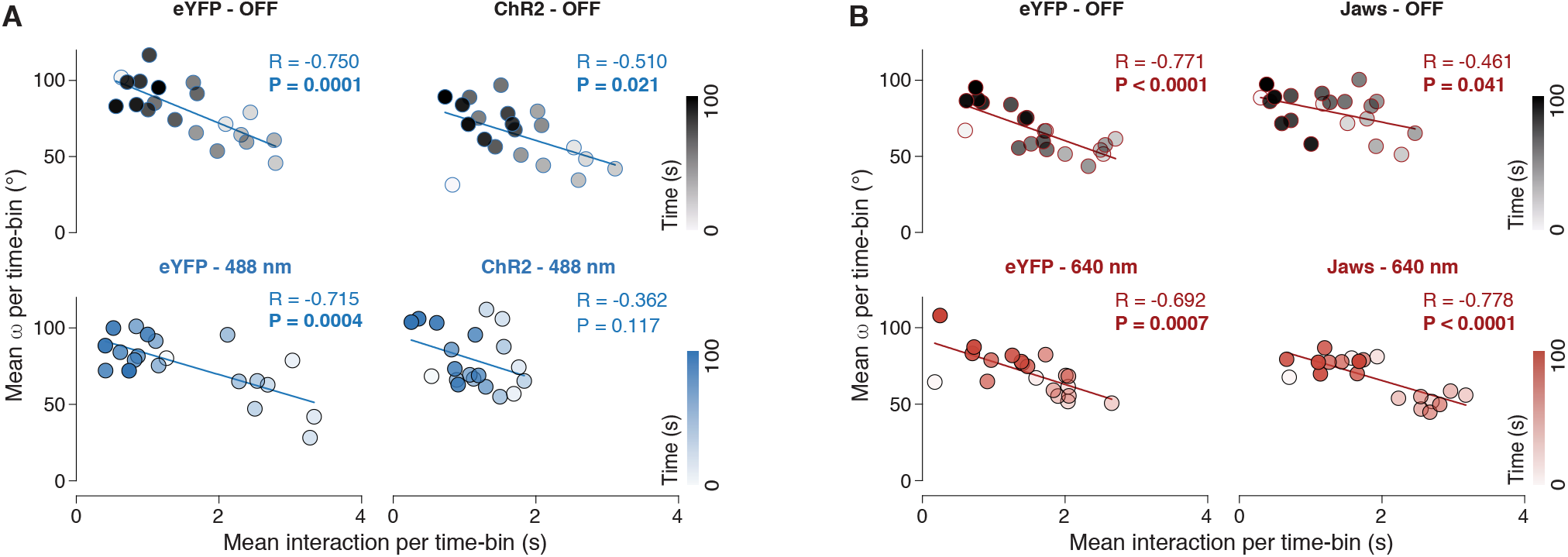
Correlation between interaction time and head-orientation angle during social interaction task. **(A-B)** Linear regressions between the mean head-oriented angle and the mean time of interaction per time-bin (= 5 seconds) across the free interaction session for eYFP-, ChR2- and Jaws-expressing mice in the SC for light OFF and light ON epochs. The time across the free interaction session is indicated by the colour of the point (colour-scale on the lower right corner of each graph). Pearson Correlation Coefficient was performed and values are indicated on the graphs.

**Supplementary figure 5:**
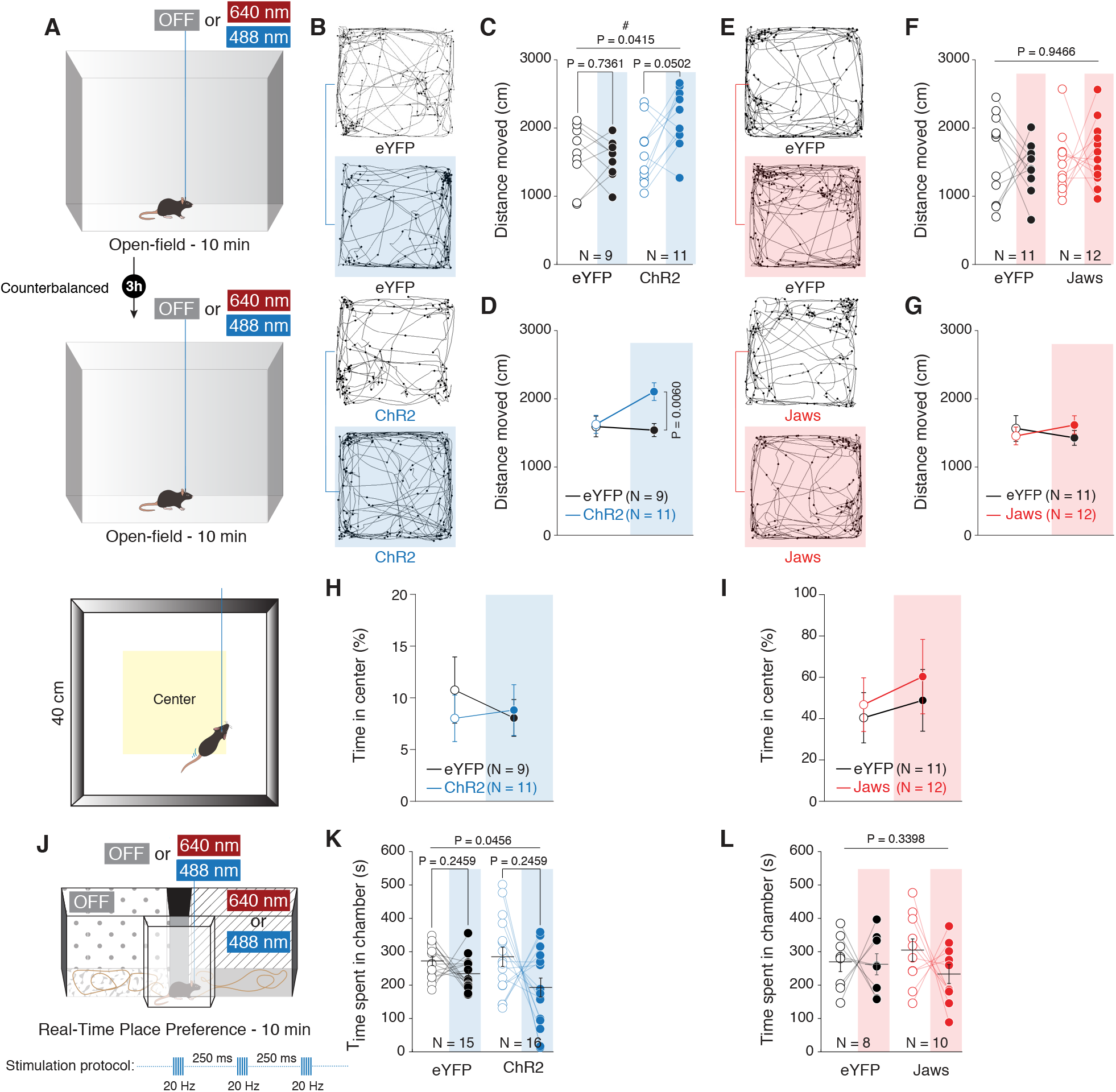
SC – VTA pathway stimulation increases exploratory behaviour and does not induce place preference. **(A)** Upper panel: schema of the open-field arena The mice are free to explore the apparatus during 10 mins in both stimulations’ conditions. Lower panel: schema of the open-field arena with the center area. **(B)** Tracking examples for eYFP and ChR2 mice in light and no-light stimulation conditions. **(C-D)** Distance moved in the open-field arena for eYFP and ChR2 mice under light and no-light stimulation. RM two-way ANOVA (Light main effect: F_(1,18)_ = 4.0046, P = 0.0607; Virus main effect: F_(1,18)_ = 4.8210, P = 0.0415; Light x Virus interaction: F_(1,18)_ = 4.8217, P = 0.0415) followed by Bonferroni-Holm post-hoc test correction. **(E)** Tracking examples for eYFP and Jaws mice in light and no-light stimulation conditions. **(F-G)** Distance moved in the open-field arena for eYFP and Jaws mice under light and no-light stimulation. RM two-way ANOVA (Light main effect: F_(1,21)_ = 0.0046, P = 0.9466; Virus main effect: F_(1,21)_ = 0.1789, P = 0.6766; Light x Virus interaction: F_(1,21)_ = 0.7033, P = 0.4111). **(H)** Time passed in the center of the open-field arena for eYFP and ChR2 mice under light and no-light stimulation. RM two-way ANOVA (Light main effect: F_(1,18)_ = 0.1628, P = 0.6913; Virus main effect: F_(1,18)_ = 0.145, P = 0.7078; Light x Virus interaction: F_(1,18)_ = 0.546, P = 0.4695). **(I)** Time passed in the center of the open-field arena for eYFP and Jaws mice under light and no-light stimulation. RM two-way ANOVA (Light main effect: F_(1,21)_ = 0.4212, P = 0.5234; Virus main effect: F_(1,21)_ = 0.5217, P = 0.4781; Light x Virus interaction: F_(1,21)_ = 0.02339, P = 0.8799). **(J)** Schema of the real-time place preference set up. The optogenetic stimulation is assigned to one chamber while the other is not associated with any stimulation. The mice are free to explore the apparatus during 10 mins. **(K)** Time spent in the chamber associated with the photostimulation or not for eYFP and ChR2 mice. RM two-way ANOVA (Light main effect: F_(1,29)_ = 4.3617, P = 0.0456; Virus main effect: F_(1,29)_ = 5.2176, P = 0.0299; Light x Virus Interaction: F_(1,29)_ = 0.7305, P = 0.3997) followed by Bonferroni-Holm post-hoc test correction. **(L)** Time spent in the chamber associated with the photostimulation or not for eYFPand Jaws mice. RM two-way ANOVA (Light main effect: F_(1,16)_ = 0.9683, P = 0.3398; Virus main effect: F_(1,16)_ = 0.2699, P = 0.6105; Light x Virus Interaction: F_(1,16)_ = 0.5421, P = 0.4722). N indicates number of mice. # indicates significantly different interaction. Error bars report s.e.m.

